# Antigenic Mapping of H2 Influenza Viruses recognized by Ferret and Human Sera and Predicting Antigenically Significant Sites

**DOI:** 10.1101/2025.11.12.687988

**Authors:** Z. Beau Reneer, Cameron Ngyen, Matthew R. Corn, Jordan L. Paugh, Owen C. Reynolds, Eric O’Hara, Ted M. Ross, Ralph S. Baric

## Abstract

Influenza viruses cause hundreds of thousands of infections globally every year. In the past century, seasonal influenza viruses have included H1N1, H2N2 and H3N2 strains. H2N2 influenza viruses circulated in the human population between 1957-1968. Previously, our group demonstrated a lack of H2N2 influenza virus immunity in individuals born after 1968, as well as the effectiveness of hemagglutinin (HA) based vaccines for multiple influenza virus subtypes. In this study, H2 antigenic maps and radial graphs were generated using previously published data from H2 HA vaccinations of ferrets and seasonal influenza vaccinations of humans. The antigenic maps revealed a stark difference in clustering of HA antigens between the ferrets and humans, and the radial graphs showed specific antigen recognition varies greatly between different influenza preimmune ferrets. These maps also revealed the significant impact that different pre-existing immunities have on antigenic recognition and clustering of antigens after vaccine boost. From these data, we predicted two possible antigenically significant sites containing various mutations that have not been previously reported and showed that one of these sites is relevant using mouse anti-sera.

**Importance:** H2N2 influenza viruses have caused at least one known pandemic in humans and are poised to cause future pandemics. Investigating the antigenic diversity of H2 HA proteins provides valuable data for designing and understanding the performance of current and future vaccines. Data evaluating the difference in antigen recognition across species and pre-existing immunity can be used to predict antigenically significant sites and evaluate the impact of H1 and H3 infection and immune imprinting on H2 vaccine immunogenicity. This information can direct future studies when both extrapolating animal data to human studies and creating next generation vaccines. Contrasting the relationships between new, contemporary and ancestral H2 HA antigens by antigenic cartography is imperative for identifying new variants of concern and updating vaccine formulations.

## Introduction

The 1957 influenza pandemic was caused by a novel H2N2 influenza virus entering the human population [1]. This novel H2N2 influenza virus was the result of a reassortment event between an avian H2N2 influenza virus and a human seasonal H1N1 virus strain [1]. The new H2N2 influenza virus encoded the PB1, HA and neuraminidase (NA) gene segments from the avian H2N2 influenza virus while the remaining five gene segments were derived from human H1N1 influenza virus [1]. The 1957 pandemic resulted in an estimated 1-2 million deaths worldwide [1]. Over the 11 years, H2N2 influenza viruses circulated as seasonal influenza viruses and underwent several antigenic changes in the HA protein before H3N2 replaced H2N2 in the human population in 1968 [1, 2]. Thereafter, human H2N2 infections have not been reported despite many avian strains showing efficient replication in primary human bronchial airway cells, foreshadowing the potential for future reemergence events [3, 4].

While H2N2 influenza viruses have not been isolated from humans since 1968, H2Nx influenza viruses have been isolated multiple times from avian species, poultry, swine and other mammals [4]. Additionally, humans are infected with other circulating HA subtypes of influenza viruses (H1N1 and H3N2) throughout their lives [1]. Previous exposure(s) to influenza viruses, and, in particular, the first influenza virus infection subtype that an individual experiences shapes the performance of future influenza virus vaccinations and infections [5–7]. Human populations have been infected with different influenza A viruses (H1N1, H2N2 and/or H3N2) over the last 80+ years [1]. The antigenic diversity of H2N2 influenza viruses combined with differences in pre-existing immunity, including the lack of pre-existing immunity in humans under the age of 55, poses enormous challenges for H2N2 vaccine design and testing [8, 9].

Antigenic cartography has provided novel insights into the antigenic relationships between seasonal influenza viruses, SARS-CoV-2 and human noroviruses, all of which evolve rapidly in response to human herd immunity [10–12]. In this study, we re-evaluated previously published data to investigate the differences in H2 HA antigen immune recognition [8, 9]. Using sera collected from ferrets, with or without pre-existing immunity to influenza A viruses, as well as sera from human subjects [8, 9], antigenic cartography was applied to map antigenic relationship based on different exposure histories, determine the cumulative effect of multiple mutations of the H2 HA proteins, and identify clusters of variants. Additionally, the antigenic and HAI results were re-analyzed to investigate the molecular basis for antigenic drift in human H2N2 viruses. We also evaluated the consequence of the order of infection/sequential infections (e.g., H1N1 and then H3N2 vs H3N2 and then H1N1) on H2 immune breadth and whether immune imprinting after single or multiple HA exposures altered H2 vaccine performance. While H3N2 alone and H3N2+H1N1 restricted H2 breadth after vaccination, we show that both H1N1 alone and H1N1+H3N2 preimmunity promotes H2 immune breadth after vaccination. In addition to previous recognized sites in the H2 HA glycoprotein, two additional amino acid residues (sites 140 and 151) were identified that potentially contribute to the HAI and antigenic cartography diversity of the H2 influenza viruses in a strain specific manner. The significance of these two antigenic sites was determined by generating wild-type (WT) and mutant virus-like particles (VLPs) containing mutations at either site 140 in the H2N2 virus in the strain Muskrat/Russia/14 or site 151 in strain Taiwan/64. Each mutant VLP was probed using antisera from H2 VLP vaccinated BALB/c mice, demonstrating a 3.5-to-6-fold difference in a residue and strain specific manner. These findings provide new insights for designing broadly cross-reactive H2 influenza vaccines in various animal species.

## Results

### Preimmune ferret antigenic variation of H2 HAs

HAI and neutralization titers in humans and ferrets are commonly used to evaluate the antigenic properties of influenza viruses. Herein, we use cartography to contrast and compare HAI and neutralization datasets in naïve and preimmune ferrets and humans, following natural infection and/or vaccination. Previously [8], female fitch ferrets were infected with sublethal doses of one of three subtypes of influenza viruses (H1N1, H2N3, or H3N2) in singularity or in combination to mimic primary infection heterogeneity seen in humans (Fig 1). The ferrets were infected with WT influenza viruses isolated from humans, with the exception of the H2N3 influenza virus strain which was isolated from chickens in 2004 (chicken/PA/04) [8].

**Figure 1:**
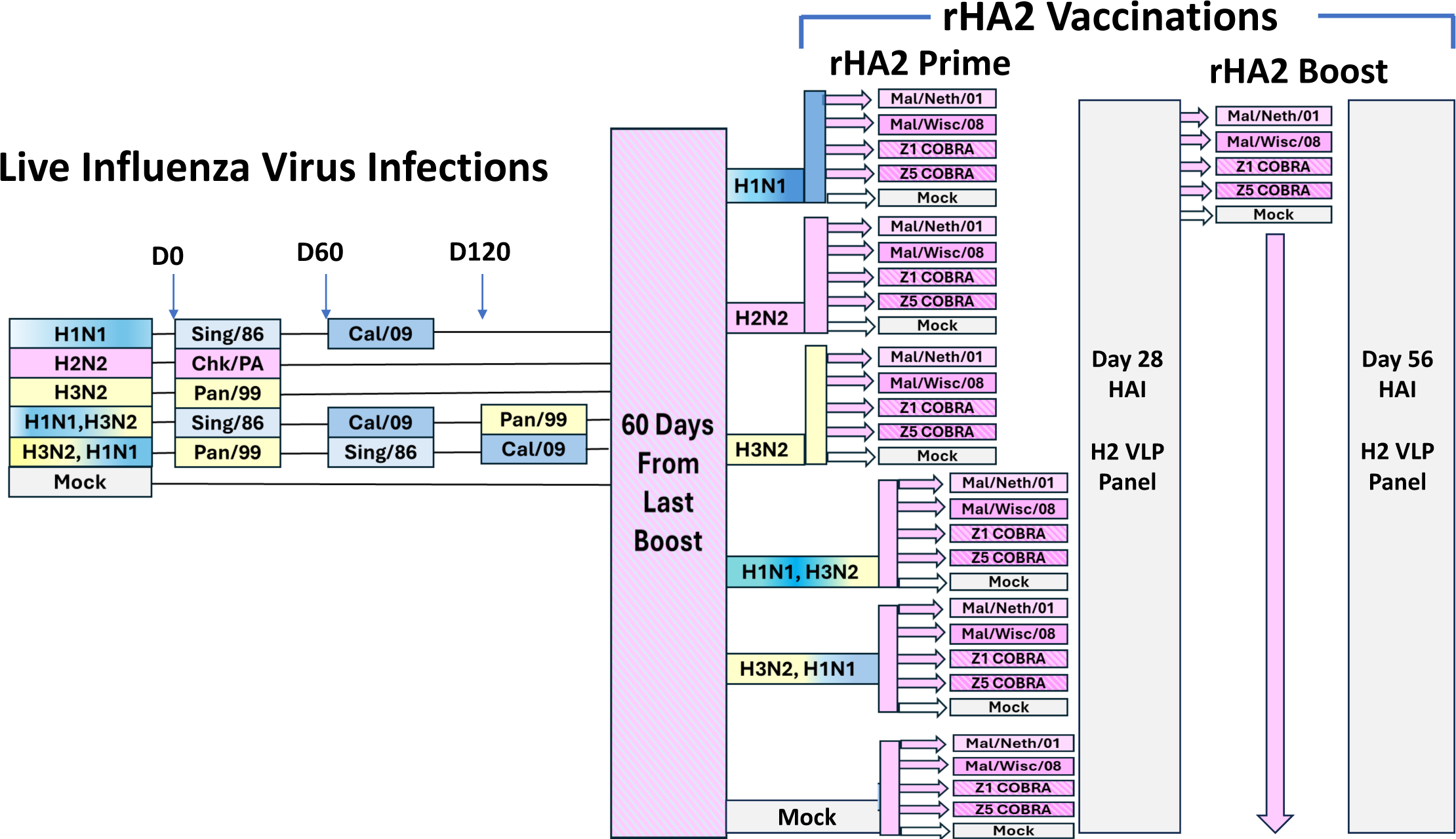
Ferret Preimmunity and Vaccination Outline. Ferrets (n=20/group) were infected with influenza viruses on D0 to establish influenza preimmunity. Ferrets were allowed to rest for 60 days before 3 groups of ferrets were infected a second time with a heterologous influenza virus. All of the ferrets were allowed to rest for 60 days before 2 of the 3 groups of ferrets were infected a third time with a heterologous influenza virus. 60 days after each ferret group’s final influenza virus infection, ferrets in each preimmune group (n=4) were vaccinated with either one of four rHA vaccines or a mock vaccines containing only PBS and adjuvant. 28 days after prime vaccination, each ferrets was vaccinated with the homologous vaccine that they received previously.

These ferrets were termed ‘preimmune’ and separated into six groups (Fig 1). The ferrets were allowed to recover for 60 days after each infection and then another 60 days before vaccination (e.g., 120 dpi). Only ferrets infected with H2N3 had Ab titers to any of the antigens in our H2 panel prior to H2 vaccinations (data not shown). The ferrets were then divided into five groups and vaccinated intramuscularly twice, 28 days apart, with 15μg of rH2 protein [*e.g.,* Mal/Neth/01, Mal/Wisc/08, COBRA Z1, COBRA Z5, mock (no antigen)] that were mixed with MF59 adjuvant (Fig 1). COBRA Z1 and Z5 antigens are candidate, broadly cross-reactive antigens that were previously designed for increased breadth and immune performance [13]. Virus-like particles (VLPs) expressing the 12 different WT H2 HAs were used for the HAI assay and live viruses were used for neutralization assays. NA antibodies elicited by the H1N1, H2N3 and H3N2 viral infections did not to contribute to HAI titers (due to assay having HA Ab specificity) and are unlikely to have contributed to neutralization titers due to mock vaccinated ferrets having neut titers at or below the limit of detection.

After infection, sera collected from ferrets infected with the H2N3 viruses elicited antibodies with robust HAI responses against the H2 VLP panel, while ferrets singly infected the H1N1 or H3N2 influenza viruses had sporadic and lower-level HAI titers against the H2 VLPs (Table 1). After primary H2 vaccination, all vaccinated H2N3 preimmune ferrets had an average HAI titer ≥1:40 against all VLPs in the panel, representing a ≥4-fold increase in HAI titer. While both the H1N1 and H3N2 preimmune, mock vaccinated ferrets had average HAI titers ≤1:5 against the entire H2 VLP panel, the H2N3 preimmune, mock vaccinated ferrets had serum HAI titers ≥1:40 against seven of the twelve strains (Table 1). Compared to COBRA Z5 and homotypic H2N3 infecting strains, COBRA Z1 vaccinated animals had nearly equivalent or higher HAI titers against most of the H2 VLPs assessed in our panel.

**Table 1:** Ferret HAls after singlee vaccination.

| Table 1: Ferret HAI<sub>s</sub> after single vaccination |  |  |  |  |  |  |  |  |  |  |  |  |  |  |
| --- | --- | --- | --- | --- | --- | --- | --- | --- | --- | --- | --- | --- | --- | --- |
| Pre-Immunity | Sera | Mal/NL/01 | Chk/Pots/84 | Musk/Rus/14 | Duk/Cam/13 | J/57 | Mosc/65 | T/64 | Duk/HK/78 | Mal/WI/08 | Sw/MO/06 | Qu/RI/16 | Tur/CA/08 | Average |
| H1N1 | Mal/Neth/01 | 94 | 8 | 83 | 5 | 10 | 188 | 25 | 9 | 25 | 35 | 203 | 40 | 60 |
| H1N1 | Mal/Wisc/08 | 126 | 96 | 124 | 5 | 29 | 170 | 41 | 25 | 93 | 93 | 261 | 143 | 101 |
| H1N1 | Z1 COBRA | 18 | 19 | 5 | 5 | 13 | 145 | 11 | 13 | 36 | 64 | 63 | 46 | 37 |
| H1N1 | Z5 COBRA | 6 | 8 | 5 | 5 | 25 | 41 | 8 | 6 | 16 | 16 | 14 | 19 | 14 |
| H1N1 | Mock | 5 | 5 | 5 | 5 | 5 | 5 | 5 | 5 | 5 | 5 | 5 | 5 | 5 |
| H2N3 | Mal/Neth/01 | 561 | 641 | 121 | 1680 | 441 | 641 | 181 | 1401 | 1921 | 2561 | 881 | 2561 | 1133 |
| H2N3 | Mal/Wisc/08 | 1000 | 1280 | 210 | 220 | 500 | 1360 | 440 | 520 | 3360 | 2720 | 1040 | 3200 | 1321 |
| H2N3 | Z1 COBRA | 1200 | 1200 | 190 | 1280 | 520 | 1360 | 400 | 680 | 2720 | 2080 | 1440 | 2240 | 1276 |
| H2N3 | Z5 COBRA | 220 | 560 | 80 | 370 | 200 | 640 | 140 | 200 | 720 | 880 | 420 | 800 | 436 |
| H2N3 | Mock | 45 | 70 | 5 | 5 | 100 | 100 | 5 | 25 | 14 | 140 | 100 | 70 | 57 |
| H3N2 | Mal/Neth/01 | 5 | 5 | 5 | 15 | 5 | 14 | 5 | 5 | 5 | 8 | 5 | 6 | 7 |
| H3N2 | Mal/Wisc/08 | 5 | 5 | 5 | 6 | 5 | 15 | 5 | 5 | 5 | 8 | 5 | 6 | 6 |
| H3N2 | Z1 COBRA | 15 | 14 | 5 | 6 | 8 | 85 | 5 | 5 | 26 | 43 | 41 | 43 | 25 |
| H3N2 | Z5 COBRA | 5 | 11 | 5 | 5 | 6 | 38 | 5 | 5 | 11 | 19 | 15 | 19 | 12 |
| H3N2 | Mock | 6 | 5 | 5 | 5 | 5 | 5 | 5 | 5 | 5 | 6 | 5 | 5 | 5 |
| None | Mal/Neth/01 | 5 | 5 | 5 | 5 | 5 | 5 | 5 | 5 | 5 | 5 | 5 | 5 | 5 |
| None | Mal/Wisc/08 | 5 | 5 | 5 | 5 | 5 | 5 | 5 | 5 | 5 | 5 | 5 | 5 | 5 |
| None | Z1 COBRA | 9 | 9 | 5 | 5 | 24 | 24 | 5 | 5 | 5 | 5 | 36 | 9 | 12 |
| None | Z5 COBRA | 5 | 5 | 5 | 5 | 5 | 5 | 5 | 5 | 5 | 5 | 5 | 5 | 5 |
| None | Mock | 5 | 5 | 5 | 5 | 5 | 5 | 5 | 5 | 5 | 5 | 5 | 5 | 5 |
| H1N1, H3N2 | Mal/Neth/01 | 86 | 91 | 48 | 646 | 25 | 123 | 15 | 28 | 15 | 183 | 106 | 181 | 129 |
| H1N1, H3N2 | Mal/Wisc/08 | 695 | 1350 | 683 | 760 | 56 | 490 | 13 | 103 | 400 | 480 | 430 | 320 | 482 |
| H1N1, H3N2 | Z1 COBRA | 348 | 390 | 328 | 1375 | 45 | 1040 | 35 | 120 | 400 | 360 | 300 | 360 | 425 |
| H1N1, H3N2 | Z5 COBRA | 23 | 55 | 9 | 686 | 18 | 190 | 11 | 24 | 50 | 140 | 88 | 100 | 116 |
| H1N1, H3N2 | Mock | 5 | 5 | 5 | 5 | 5 | 5 | 5 | 5 | 5 | 5 | 5 | 5 | 5 |
| H3N2, H1N1 | Mal/Neth/01 | 5 | 5 | 5 | 14 | 6 | 5 | 5 | 5 | 5 | 5 | 5 | 5 | 6 |
| H3N2, H1N1 | Mal/Wisc/08 | 5 | 5 | 5 | 71 | 6 | 5 | 5 | 5 | 5 | 5 | 5 | 5 | 11 |
| H3N2, H1N1 | Z1 COBRA | 5 | 5 | 5 | 5 | 6 | 5 | 5 | 5 | 5 | 5 | 5 | 5 | 5 |
| H3N2, H1N1 | Z5 COBRA | 5 | 5 | 5 | 41 | 6 | 5 | 5 | 5 | 5 | 5 | 5 | 5 | 8 |
| H3N2, H1N1 | Mock | 5 | 5 | 5 | 5 | 5 | 5 | 5 | 5 | 5 | 5 | 5 | 5 | 5 |
| Average |  | 151 | 196 | 66 | 242 | 70 | 224 | 47 | 108 | 329 | 330 | 184 | 341 | 191 |

Excluding the mock vaccinated ferrets, each of the H1N1 preimmune ferret vaccine groups had similar or higher HAI titer averages to each H2 antigen in our panel than the H3N2 preimmune ferrets. Ferrets preimmune to the H3N2 influenza virus had low HAI titers against each of the H2 VLPs in the panel and no ferret had a >4-fold increase in HAI activity against the H2 influenza viruses compared to mock controls after vaccination (HAI≤5). However, both COBRA Z1 and Z5 vaccinated ferrets had a >4-fold increase in HAI titer against 2-4 of the twelve H2 VLPs in the panel (Table 1). While none of the mock vaccinated ferrets had detectable HAI titers against any H2 VLP, two ferrets had preimmune titers to H1N1 influenza virus (average HAI titer ≥1:40 against the Mosc/65 VLP) and 11/12 ferrets had a ≥4-fold increase in HAI titers HAI against the Mal/WI/08 and 8/12 against the Mal/NL/01 over background (Table 1). HAI titers in the Z1 and Z5 COBRA vaccinated ferrets had lower responses than ferrets vaccinated with H2 Mal/WI/08 (5/12) and Mal/NL/01 (2/12) with a ≥4-fold increased HAI titer over mock controls (Table 1).

Additional groups of ferrets were primed with sequential infections of two influenza viruses, either H1N1 followed by H3N2 or H3N2 followed by H1N1. The mock vaccinated ferrets had HAI titers of ≤1:5 across the H2 panel in both of the double infected, preimmune group. Across all H2 vaccine groups, ferrets that were preimmune after H1N1 followed by H3N2 influenza virus infections, demonstrated an average HAI >1:40 against 8 to 11 of the 12 VLPs in our panel. While HAI titers elicited by the Z5 COBRA rHA vaccine were generally less robust across the HAI panel, ferrets vaccinated with the Z1 COBRA HA vaccine had a ≥4-fold increase in neutralization titers against all 12 of the H2 VLPs (Table 1). In the H3N2 and H1N1 preimmune group, H2 HAI titers were severely attenuated across the panel. In fact, the Mal/WI/08 and Z5 COBRA vaccinated ferrets only had a ≥4-fold increase in neutralization titers against a single H2 VLP (Duke/Cam/13) (Table 1), showing the significance of influenza A virus infection order and immune imprinting on downstream immune responses to HA2 antigens.

### Antigenic Relationships across the Preimmune Groups by Cartography

Radial graphs were generated to clearly visualize the impact of preexposure histories and vaccines on antigen recognition (Fig 2). On D14, the H2 preimmune ferrets had the highest HAI titers and recognized the highest number of antigens of any preimmune group. The mock preimmune group had the lowest average magnitude HAI titers (Fig 2), while the H3, H1 preimmune ferrets only had HAI titers to Duk/Cam/13. Being infected with H3N2 alone or prior to an H1N1 infection dramatically impacted H2 vaccine performance in ferrets.

**Figure 2.**
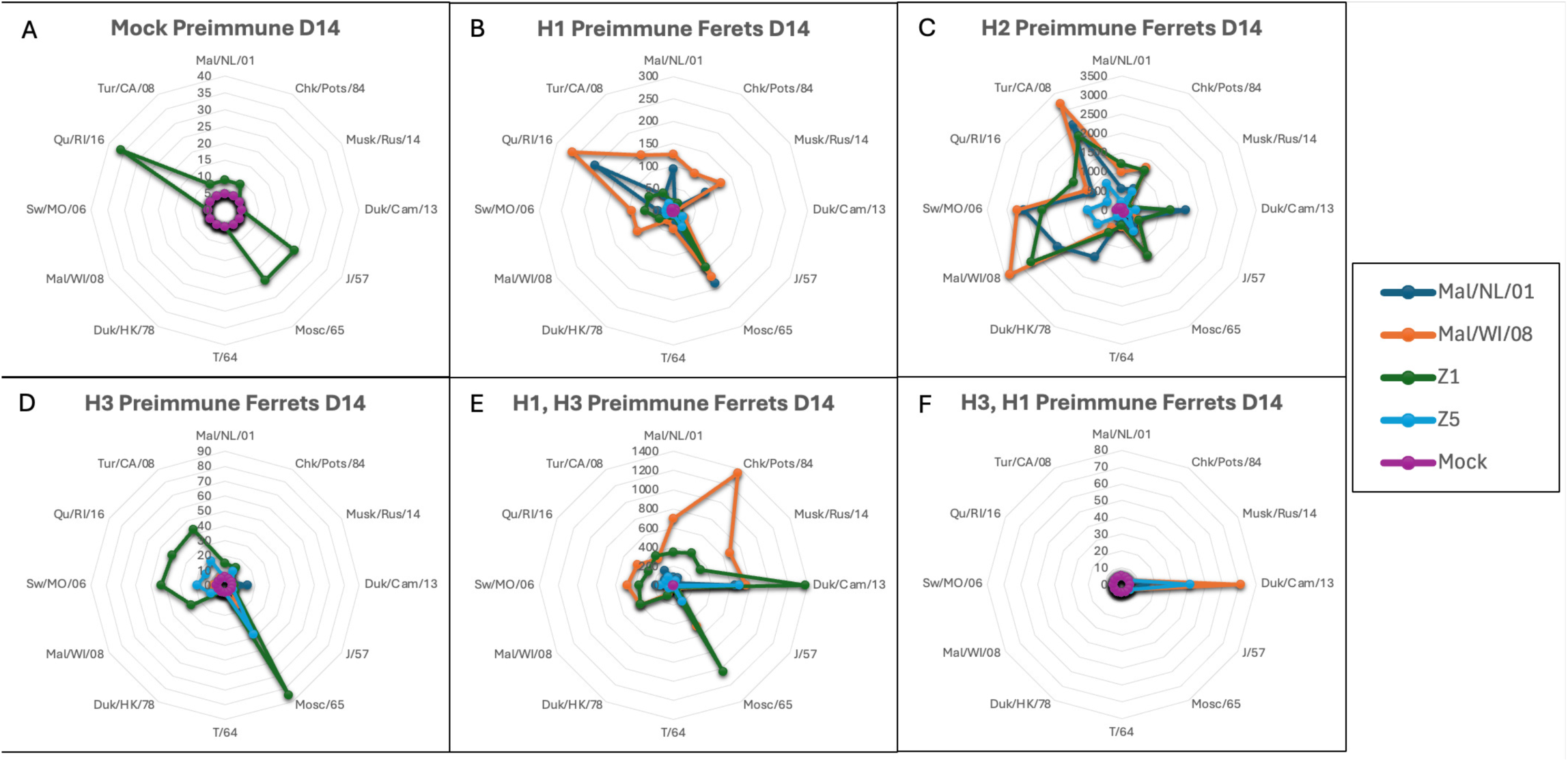
HAI radial graph post-prime H2 vaccination. Vaccine groups are colored coated per the legend and consistent between each graph. The circles in each graph represent the HAI titer. (A) Mock preimmune ferrets, (B) H1 Preimmune ferrets, (C) H2 Preimmune ferrets, (D) H3 Preimmune ferrets, (E) H1, H3 Preimmune ferrets, (F) H3, H1 Preimmune ferrets.

Antigenic cartography was applied to further examine the HAI data following the first vaccinations or infections (Fig 3). In antigenic cartography, antigens and sera that cluster in the center respond most similarly while those that radiate outward are increasingly divergent from the other antigens/sera. Each of the antigens clustered together near the center of the graph with the exception of the Duk/Cam/13 HA antigen (Fig 3a-b). In general, the sera from each preimmune group clustered together near the center of the graph, which was especially evident in the H2N3 and H3N2 vaccinated ferrets. For the other preimmune groups, ferrets preimmune to the H1N1 influenza virus, and to a greater extent the H1N1-H3N2 and the H3N2-H1N1 preimmune ferrets, had a significantly altered serological response with expanded neutralization breadth against the H2 VLP panel (Fig 3a-b). The serum samples collected from the four naïve preimmune groups all overlapped on the antigenic graph since there was no variability in their HAI titers (Fig 3a-b).

**Figure 3.**
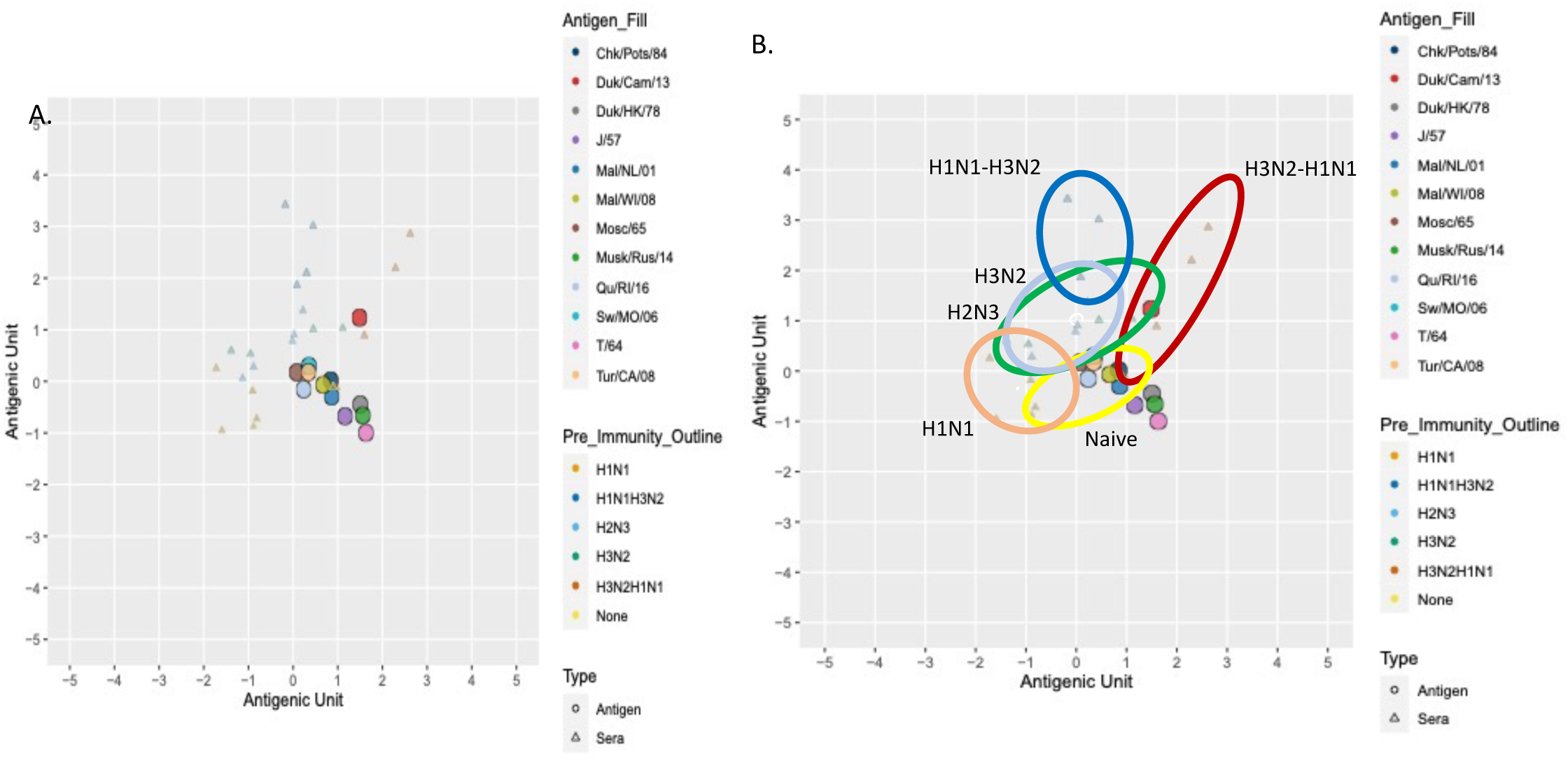
Ferret HAI Antigenic Cartography Maps After Prime Vaccination. (A) Antigenic maps were generated for each ferret preimmune group in HAIs. Antigens are represented as circles and sera from the ferrets are represented by triangles. (B) The sera from each influenza preimmune groups are grouped within each colored circle for clarity.

### Serum HAI Responses after Secondary Vaccinations

Preimmune ferrets were boosted with the same H2 antigen and convalescent serum was isolated on day 56 post-boost (Fig 1). After the second vaccination, the H2N3 preimmune ferret HAI titers did not significantly change compared to the HAI titers following the first vaccination (Tables 1, 2). Post boost, in ferrets preimmune after H1N1 influenza virus infection, the H2 HAI titers not only increased slightly against the Mal/WI/08 and MAL/NL/01 VLPs but a broader ≥4-fold increase in H2 HAI titers was noted against the other H2 strains. However, in the COBRA vaccine groups, the mean H2 HAI titers increased ≥7-10 fold over preimmune vaccinated animals demonstrating their superiority as vaccine antigens compared to WT antigens (Table 2). In the non-preimmune group, each of the rH2 vaccinated ferrets showed increases in HAI titers after boost (Table 2). The Mal/NL/01 and Mal/WI/08 vaccination groups had average HAI titers >1:40 with four-fold elevated titers in 8/12 and 7/12 antigens, respectively (Table 2). The Z1 and Z5 COBRA vaccination groups had an average HAI >1:40 to eleven and nine of the twelve of the VLPs in the panel with 11/12 and 10/12 4-fold elevated titers over baseline, respectively (Table 2). In the H3N2 preimmune group, the Mal/NL/01 and Mal/WI/08 vaccination groups had, on average, HAI >1:40 to seven and eleven of the twelve VLPS in the panel (Table 2). Both the Z1 and Z5 COBRA vaccine groups had, on average, HAI >1:40 to ten of the twelve VLPs in the panel (Table 2). Overall, the H2N3 and H1N1 preimmune ferrets had higher HAI titers to the H2 VLPs than the H3N2 preimmune ferrets.

**Table 2:** Ferret HAI after second vaccination.

| Table 2: Ferret HAI after second vaccination |  |  |  |  |  |  |  |  |  |  |  |  |  |  |
| --- | --- | --- | --- | --- | --- | --- | --- | --- | --- | --- | --- | --- | --- | --- |
| Pre-Immunity | Sera | Mal/NL/01 | Chk/Pots/84 | Musk/Rus/14 | Duk/Cam/13 | J/57 | Mosc/65 | T/64 | Duk/HK/78 | Mal/WI/08 | Sw/MO/06 | Qu/RI/16 | Tur/CA/08 | Average |
| H1N1 | Mal/Neth/01 | 85 | 60 | 124 | 5 | 55 | 280 | 49 | 40 | 180 | 180 | 360 | 320 | 145 |
| H1N1 | Mal/Wisc/08 | 83 | 23 | 6 | 5 | 53 | 300 | 13 | 33 | 110 | 90 | 220 | 220 | 96 |
| H1N1 | Z1 COBRA | 130 | 113 | 35 | 5 | 111 | 885 | 58 | 111 | 450 | 225 | 500 | 450 | 256 |
| H1N1 | Z5 COBRA | 135 | 80 | 88 | 5 | 63 | 260 | 165 | 43 | 220 | 290 | 530 | 340 | 185 |
| H1N1 | Mock | 5 | 5 | 5 | 5 | 5 | 5 | 5 | 5 | 5 | 5 | 5 | 5 | 5 |
| H2N3 | Mal/Neth/01 | 521 | 201 | 101 | 521 | 261 | 561 | 121 | 181 | 1121 | 1281 | 281 | 1121 | 523 |
| H2N3 | Mal/Wisc/08 | 1060 | 480 | 120 | 520 | 380 | 800 | 220 | 220 | 1680 | 2720 | 760 | 1760 | 893 |
| H2N3 | Z1 COBRA | 1840 | 880 | 200 | 1840 | 440 | 1600 | 380 | 440 | 2080 | 2080 | 1040 | 2080 | 1242 |
| H2N3 | Z5 COBRA | 340 | 480 | 120 | 260 | 150 | 360 | 120 | 160 | 720 | 760 | 420 | 560 | 371 |
| H2N3 | Mock | 45 | 70 | 5 | 5 | 100 | 100 | 5 | 25 | 14 | 140 | 100 | 70 | 57 |
| H3N2 | Mal/Neth/01 | 28 | 41 | 5 | 966 | 15 | 131 | 5 | 26 | 51 | 51 | 56 | 56 | 119 |
| H3N2 | Mal/Wisc/08 | 70 | 70 | 5 | 1284 | 50 | 280 | 45 | 50 | 260 | 360 | 180 | 360 | 251 |
| H3N2 | Z1 COBRA | 85 | 130 | 10 | 1363 | 83 | 340 | 19 | 73 | 300 | 600 | 340 | 680 | 335 |
| H3N2 | Z5 COBRA | 105 | 90 | 6 | 1481 | 85 | 360 | 9 | 85 | 180 | 320 | 300 | 340 | 280 |
| H3N2 | Mock | 5 | 5 | 5 | 5 | 5 | 5 | 5 | 5 | 5 | 5 | 5 | 5 | 5 |
| None | Mal/Neth/01 | 25 | 25 | 5 | 33 | 24 | 83 | 5 | 5 | 5 | 63 | 80 | 23 | 31 |
| None | Mal/Wisc/08 | 36 | 26 | 5 | 5 | 83 | 163 | 5 | 5 | 5 | 93 | 81 | 41 | 46 |
| None | Z1 COBRA | 320 | 160 | 5 | 400 | 600 | 600 | 44 | 121 | 320 | 640 | 340 | 210 | 313 |
| None | Z5 COBRA | 48 | 66 | 5 | 84 | 241 | 141 | 5 | 24 | 44 | 141 | 115 | 65 | 82 |
| None | Mock | 5 | 5 | 5 | 5 | 5 | 5 | 5 | 5 | 5 | 5 | 5 | 5 | 5 |
| H1N1, H3N2 | Mal/Neth/01 | 205 | 185 | 94 | 173 | 48 | 243 | 28 | 48 | 183 | 363 | 351 | 183 | 175 |
| H1N1, H3N2 | Mal/Wisc/08 | 480 | 420 | 130 | 980 | 85 | 500 | 28 | 63 | 480 | 1600 | 1480 | 440 | 557 |
| H1N1, H3N2 | Z1 COBRA | 280 | 320 | 115 | 280 | 120 | 800 | 60 | 100 | 320 | 800 | 440 | 280 | 326 |
| H1N1, H3N2 | Z5 COBRA | 130 | 100 | 25 | 83 | 55 | 220 | 20 | 38 | 130 | 190 | 150 | 110 | 104 |
| H1N1, H3N2 | Mock | 5 | 5 | 5 | 5 | 5 | 5 | 5 | 5 | 5 | 5 | 5 | 5 | 5 |
| H3N2, H1N1 | Mal/Neth/01 | 34 | 83 | 181 | 600 | 360 | 84 | 71 | 5 | 29 | 201 | 53 | 61 | 147 |
| H3N2, H1N1 | Mal/Wisc/08 | 21 | 94 | 241 | 580 | 210 | 48 | 41 | 5 | 63 | 300 | 104 | 141 | 154 |
| H3N2, H1N1 | Z1 COBRA | 24 | 33 | 250 | 600 | 290 | 25 | 44 | 5 | 25 | 330 | 6 | 86 | 143 |
| H3N2, H1N1 | Z5 COBRA | 21 | 31 | 105 | 290 | 166 | 26 | 26 | 5 | 28 | 105 | 33 | 55 | 74 |
| H3N2, H1N1 | Mock | 5 | 5 | 5 | 5 | 5 | 5 | 5 | 5 | 5 | 5 | 5 | 5 | 5 |
| Average |  | 206 | 143 | 67 | 413 | 138 | 307 | 54 | 65 | 301 | 465 | 278 | 336 | 231 |

In the H1N1, H3N2 preimmune group, animals responded more efficiently against the H2 virus after vaccine boost than similarly boosted H1N1 and H3N2 preimmune groups alone. For example, H1N1, H3N2 preimmune animals mounted ≥4-fold increased H2 HAI responses to 11/12 H2 VLPs in Mal/NL/01 and Mal/WI/08 vaccination groups, with average titers of ∼1/129 and 1/482, respectively. Again, the Z1 COBRA vaccine proved superior, as ≥4-fold increased H2 VLP titers were evident over mocks, and increased mean titers of 1/425 over 1/129, respectively. Both titers were superior to Z5 COBRA responses as well (Table 2). Surprisingly, the H3N2, H1N1 preimmune group significantly attenuated overall H2 HAI titers in all H2 infection and vaccine groups as compared with H1N1, H3N2 vaccination. For example, the Mal/WI/08 and Z5 COBRA groups showed a ≥4-fold increased HAI response over mocks, against a single H2 VLP (Table 2). strain vaccination groups had, on average, HAI >1:40 against nine and ten of the twelve VLPs in the panel, respectively (Table 2). However, both the Z1 COBRA and Z5 COBRA vaccine groups had, on average, HAI >1:40 to six and five of the twelve VLPs in the panel respectively (Table 2). It is noteworthy that H3N1, H1N1 preimmune animals mounted minimal responses against Dck/HK/78.

### Radial Graphs and Cartography after H2 Vaccine Boost

Radial graphs were generated to visualize the HAI responses in the preimmune ferrets after H2 boost (Fig 4). Overall, the HAI titers increased to all antigens except for the Mal/WI/08 vaccinated, H1, H3 preimmune ferrets who showed a dramatic decrease in HAI titers to the Chk/Pots/84 antigen (Fig 4). The H3, H1 preimmune ferret and the H3 preimmune ferret HAI responses predominately increased only to Duk/Cam/13 and Duk/Cam/13 respectively (Fig 4).

**Figure 4.**
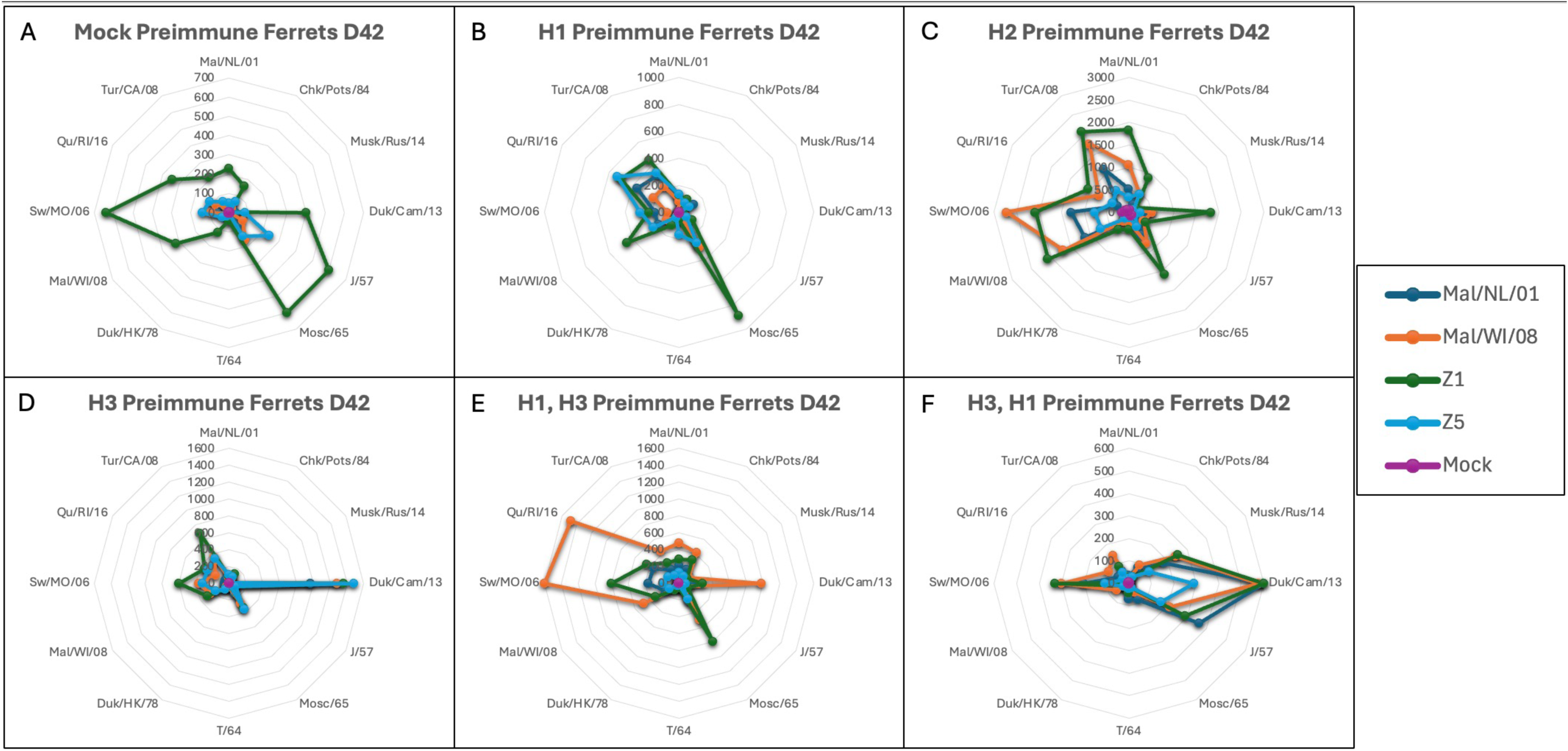
HAI radial graph post-prime H2 vaccination. Vaccine groups are colored coated per the legend and consistent between each graph. The circles in each graph represent the HAI titer. (A) Mock preimmune ferrets, (B) H1 Preimmune ferrets, (C) H2 Preimmune ferrets, (D) H3 Preimmune ferrets, (E) H1, H3 Preimmune ferrets, (F) H3, H1 Preimmune ferrets.

Antigenic cartography was used to analyze HAI activity of ferret serum samples following the second vaccination (Fig 5). The HA antigens Musk/Rus/14, Duk/Cam/13, T/64 and Duk/HK/78 were antigenically farthest away from the other HA antigens (Fig 5). Sera from the preimmune groups separated from each other (Fig 5). All the serum samples collected from the naïve ferrets, as well as ferrets preimmune to H2N3, H1N1-H3N2 or H3N2-H1N1 influenza viruses were clustered closely together on the antigenic map with the other serum samples within their same influenza group (Fig 5). Each of the preimmune animals elicited HAI responses that grouped tightly together with the other animals in its preimmune group. The H3N2 and H3N2-H1N1 preimmune ferrets had sera with the furthest antigenic distance from the other preimmune groups (Fig 5). Interesting, first exposure to H1N1 appeared to elicit imprinted responses that were more favorable to H2N2 vaccination, as compared to H3N2 first exposure.

**Figure 5.**
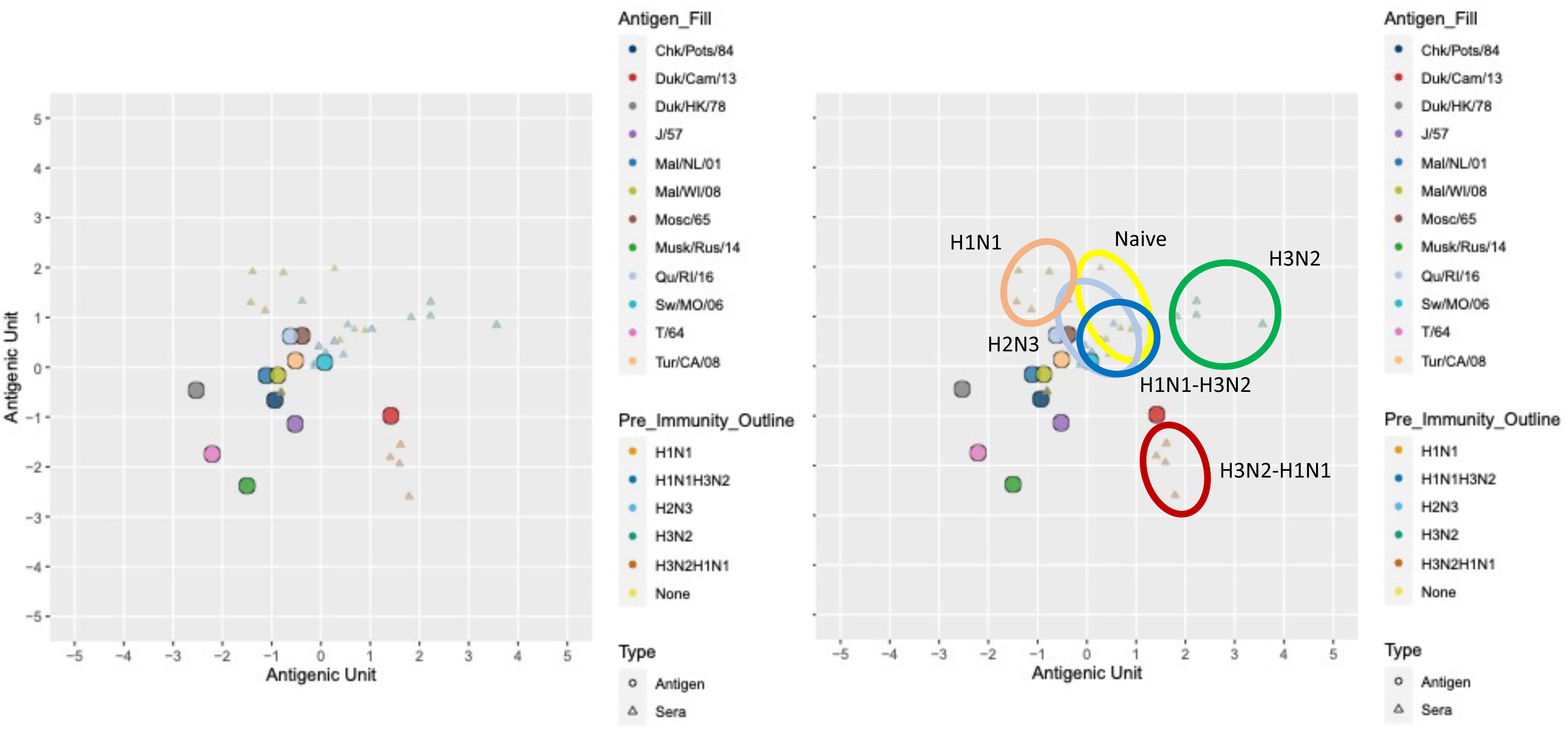
Ferret HAI Antigenic Cartography Maps After Boost Vaccination. (A) Antigenic maps were generated for each ferret preimmune group in HAIs. Antigens are represented as circles and sera from the ferrets are represented by triangles. The preimmune groups are beginning to separate from each other and the antigens due to increased HAI titer differences compared to prime vaccinations. (B) The sera from each influenza preimmune groups are grouped within each colored circle for clarity.

Based on the antigenic cartography, the Musk/Rus/14, T/64 and Duk/HK/78 sera were separated from the other sera in the HAI antigenic map (Fig 5). T/64 has mutations at sites 132, 139 and 188. Linster et al. found that site 139 (Asparagine [N] to Lysine [K]) had the most substantial antigenic change of any of the single amino acid mutations [2] (Table S1). Both T/64 and Musk/Rus/14 had the N to K mutation which is likely driving their divergence from the other sera and antigens respectively (Table S1). Mutations at sites 132 and 184 both had minor changes in antigenic distance individually. Both Musk/Rus/14 and T/64 have a Glutamine (Q) and Methionine (M) respectively at site 132 while all the other viruses (except Chik/Pots/84) encode Arginine (R) (Table S1).

Additionally, both Musk/Rus/14 and T/64 have a positively charged Lysine (K) at site 139 while all other viruses encode an Asparagine (N) (Table S2). Of the sites identified by Linster et al., Duk/HK/78 did not have a significant mutation in any site [2] (Table S1). These data suggest that other amino acid residue substitutions are also contributing to the antigenic variation of Musk/Rus/14, T/64 and Duk/HK/78.

Additionally, Musk/Rus/14 has a unique mutation at site 140 which was not found by Linster et al. [2] (Table S2). This Proline [P] to Serine [S] mutation at site 140 along with the mutations at sites 132 and 139 are all likely contributing to the low HAI titers against Musk/Rus/14. Both T/64 and Duk/HK/78 also had significant mutations at site 151 which was not reported by Linster et al [2] (Table S2). T/64 encodes a Glutamic acid [E] while Duk/HK/78 encodes a Threonine [T] at site 151 while all other H2 HA encode a Lysine [K] at site 151. These mutations may contribute to the lower HAI titers against both T/64 and Duk/HK/78. In addition to site 151, Duk/HK/78 also has nine other amino acid changes that are not present in any of the other H2 influenza viruses.

To evaluate the significance of site 140 in Musk/Rus/14 and site 151 in T/64, we generated virus-like particles (VLPs) containing each mutant HA protein. We generated both the WT HA sequences of both Musk/Rus/14 and T/64 while also utilizing site-directed mutagenesis (SDM) to mutate site 140 in Musk/Rus/14 from Serine [S] to Proline [P] (S140P) and site 151 in T/64 from Glutamic acid [E] to Lysine [K] (E151K). We then vaccinated BALB/c mice with 3ug of recombinant HA (rHA) with MF59 oil-water emulsion adjuvant. The mice (n=4/group) were vaccinated twice, four weeks apart before being bled two-weeks post-boost. The antigens used for vaccinations were Z1 COBRA and Mal/NL/01 that we have used previously [13]. The average HAI titers against T/64 did increase for both Mal/NL/01 and Z1 COBRA but did not achieve statistical significance (p=0.1265 NL and p=0.2709 Z1) (Fig 6). The HAI titers against Musk/Rus/14 did not increase and remained below the limit of detection (Fig 6).

**Figure 6.**
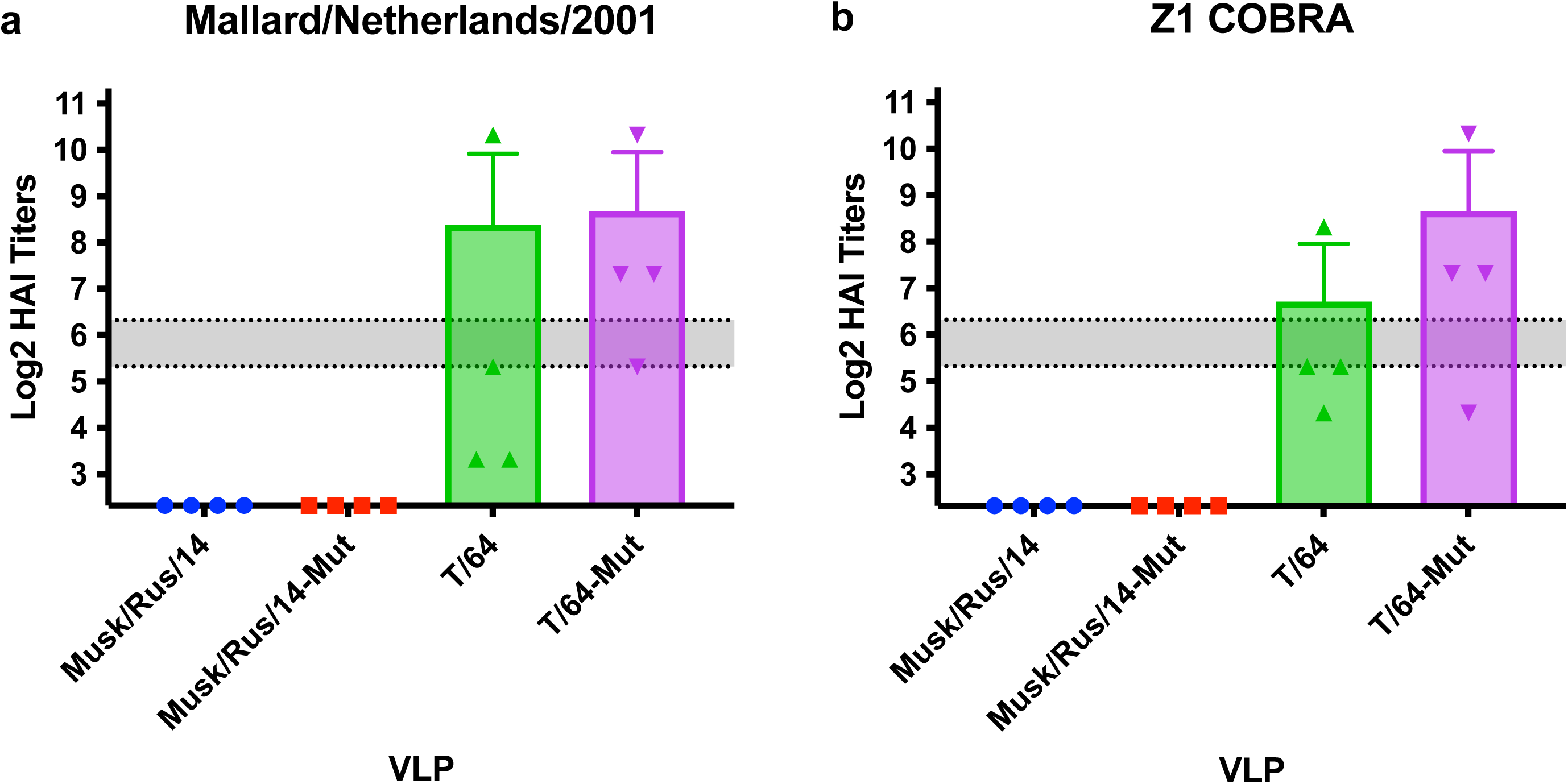
Antigenic significance of sites 140 and 151. BALB/c mice were vaccinated with 3ug of recombinant proteins (Mal/NL/01 or Z1 COBRA) with MF50 oil-water-emulsion adjuvant. (A) Mal/NL/01 vaccinated mice had HAI below the limit of detection for both the WT and Mutant (S140P) Musk/Rus/14 VLPs. The average HAI titer against the T/64 mutant (E151K) increased but was not significant over the WT T/64 VLP. (B) Z1 COBRA vaccinated mice had HAI below the limit of detection for both the WT and Mutant (S140P) Musk/Rus/14 VLPs. The average HAI titer against the T/64 mutant (E151K) increased but was not significant over the WT T/64 VLP.

### Live virus Neutralization Assays

Pooled sera collected from the ferrets after the second vaccination was used in classical neutralization assays against a panel of heterologous H2Nx influenza viruses. Neutralization assays quantify all antibodies that block viral attachment, replication and spread. The mock vaccinated ferrets all had neut titers <1:10 in all the preimmune groups apart from the H2N3 preimmune group (Table 3). In the H2N3 preimmune group, the mock vaccinated ferrets had, on average, neut titers ≥1:160 to one of the seven H2 viruses in the panel.

**Table 3:** Ferret Neutralization Titers.

| Table 3: Ferret Neutralization Titers |  |  |  |  |  |  |  |  |  |
| --- | --- | --- | --- | --- | --- | --- | --- | --- | --- |
| Pre-Immunity | Vaccine | Chk/Pots/84 | Chk/PA/04 | For/57 | T/64 | Duk/HK/78 | Sw/MO/06 | Mal/MN/08 | Averages |
| H2N2 | Mal/NL/01 | 640 | 380 | 48 | 34 | 640 | 640 | 452 | 405 |
| H2N2 | Mal/WI/08 | 640 | 537 | 48 | 56 | 640 | 640 | 640 | 457 |
| H2N2 | Z1 | 640 | 640 | 134 | 67 | 640 | 640 | 640 | 486 |
| H2N2 | Z5 | 452 | 640 | 40 | 380 | 537 | 640 | 320 | 430 |
| H2N2 | Mock | 95 | 156 | 8 | 24 | 48 | 537 | 95 | 138 |
| H3N2 | Mal/NL/01 | 17 | 7 | 5 | 20 | 12 | 452 | 17 | 76 |
| H3N2 | Mal/WI/08 | 48 | 67 | 5 | 28 | 24 | 640 | 67 | 126 |
| H3N2 | Z1 | 190 | 56 | 7 | 40 | 67 | 640 | 95 | 156 |
| H3N2 | Z5 | 113 | 113 | 5 | 20 | 34 | 640 | 48 | 139 |
| H3N2 | Mock | 5 | 7 | 5 | 5 | 5 | 6 | 6 | 6 |
| H1N1 | Mal/NL/01 | 113 | 20 | 5 | 95 | 28 | 537 | 24 | 117 |
| H1N1 | Mal/WI/08 | 113 | 40 | 6 | 537 | 34 | 640 | 226 | 228 |
| H1N1 | Z1 | 537 | 269 | 20 | 80 | 269 | 640 | 269 | 298 |
| H1N1 | Z5 | 134 | 48 | 10 | 20 | 80 | 640 | 12 | 135 |
| H1N1 | Mock | 8 | 6 | 5 | 5 | 5 | 5 | 5 | 6 |
| H3N2, H1N1 | Mal/NL/01 | 269 | 160 | 28 | 34 | 95 | 640 | 134 | 194 |
| H3N2, H1N1 | Mal/WI/08 | 113 | 113 | 10 | 34 | 67 | 640 | 134 | 159 |
| H3N2, H1N1 | Z1 | 640 | 380 | 28 | 160 | 537 | 640 | 226 | 373 |
| H3N2, H1N1 | Z5 | 226 | 160 | 17 | 48 | 160 | 640 | 160 | 202 |
| H3N2, H1N1 | Mock | 5 | 5 | 7 | 5 | 5 | 6 | 6 | 6 |
| H1N1, H3N2 | Mal/NL/01 | 640 | 640 | 34 | 5 | 80 | 640 | 640 | 383 |
| H1N1, H3N2 | Mal/WI/08 | 537 | 269 | 40 | 40 | 80 | 640 | 380 | 284 |
| H1N1, H3N2 | Z1 | 226 | 640 | 67 | 113 | 95 | 640 | 537 | 331 |
| H1N1, H3N2 | Z5 | 269 | 269 | 20 | 40 | 113 | 640 | 190 | 220 |
| H1N1, H3N2 | Mock | 5 | 6 | 5 | 5 | 5 | 5 | 5 | 5 |
| Mock | Mal/NL/01 | 34 | 14 | 5 | 10 | 10 | 640 | 20 | 105 |
| Mock | Mal/WI/08 | 12 | 17 | 5 | 5 | 7 | 640 | 20 | 101 |
| Mock | Z1 | 537 | 190 | 40 | 80 | 226 | 640 | 80 | 256 |
| Mock | Z5 | 160 | 28 | 6 | 10 | 56 | 380 | 14 | 93 |
| Mock | Mock | 5 | 5 | 5 | 5 | 5 | 5 | 5 | 5 |
| Averages |  | 247 | 196 | 22 | 67 | 153 | 512 | 182 | 197 |

Preexposure to H2N3 viruses significantly boosted immune responses across the panel. The Mal/NL/01, Mal/WI/08 and Z1 COBRA vaccination groups all had, on average, 4-fold elevated neut titers over baseline to 5/8 H2Nx viruses compared with 6/8 H2Nx viruses for Z1 COBRA. In general, neut titers ≥1:160 to five of the seven H2 viruses, while Z5 COBRA vaccination group had, on average, neut titer ≥1:160 to six of the seven H2 viruses (Table 3). In the H1N1 preimmune group, neutralization titers were reduced as compared to preimmune H2N3 vaccinated groups although the COBRA vaccines elicited 4-fold increased neut titers against more H2 antigens in the panel. Both the Mal/NL/01 and Z5 COBRA vaccination groups had, on average, a neut titer ≥1:160 to one of the seven H2 viruses while the Mal/WI/08 and Z1 COBRA vaccination groups had, on average, a neut titer ≥1:160 to three and five of the seven H2 viruses, respectively (Table 3). In the H3N2 preimmune group, the Mal/NL/01, Mal/WI/08 and Z5 COBRA vaccination groups all had, on average, a neut titer ≥1:160 to one of the seven H2 viruses, while the Z1 COBRA vaccination group had an average neut titer ≥1:160 to two of the seven H2 viruses in the panel (Table 3).

In the H1N1, H3N2 preimmune group, the Mal/NL/01, Mal/WI/08 and Z1 COBRA and Z5 COBRA vaccination groups all had an average neut titer ≥1:160 to four of the seven viruses in the panel (Table 3). In the H3N2, H1N1 preimmune group, the Mal/NL/01 vaccination group had, on average, a neut titer ≥1:160 to three of the seven H2 viruses in the panel, while the Mal/WI/08 vaccination group had, on average, a neut titer ≥1:160 to one of the seven H2 viruses in the panel. The Z1 and Z5 COBRA vaccination group had an average neut titer ≥1:160 to six and five of the seven H2 viruses in the panel, respectively (Table 3). In the non-preimmune group, each of the vaccination groups other than the mock vaccination group recognized the Sw/MO/06 virus had an average neut titer ≥1:160 (upper limit of detection). The Mal/NL/01 and Mal/WI/08 did not have an average neut titer ≥1:160 to any of the other six H2 viruses in the panel while the Z1 and Z5 COBRA vaccination groups had an average neut titer ≥1:160 to four and two of the other six H2 viruses in the panel, respectively (Table 3).

Overall, the H2N3 preimmune groups has the highest average neut titers across all seven viruses while the H3N2 and Mock preimmune groups had the lowest average neut titers across all seven viruses. Unsurprisingly, the H2N3 preimmune group had the highest average neut titers. However, the H1N1, H3N2 preimmune group also elicited superior neutralization titers as compared to the H3N2, H1N1 preimmune groups across all seven viruses with an average neut titer ≥1:220 for all of the non-mock vaccinated groups. The Z1 COBRA vaccination group also had the highest average neut titer across all six preimmune vaccine groups.

### Antigenic Cartography: Live virus neutralization assays

The antigenic cartography for the ferret neutralization data showed that the Chk/Pots/84, Chk/PA/04, Duk/HK/78 and Mal/MN/08 antigens clustered together near the center of map with the Sw/MO/06, For/57 and T/64 antigens all separated from each other (Fig. 7). The sera from the various preimmune ferret groups had significant overlap with most sera clustering near most of the antigens at the center of the map. The sera were closest to the Sw/MO/06 antigen, while being furthest from the For/57 and T/64 antigens (Fig 7). The For/57 virus did not have a significant mutation in any site identified by Linster et al [2]. Much like the HAI data from the Duk/HK/78 virus, the For/57 data indicates that are likely other amino acid sites responsible for the antigenic variation.

**Figure 7.**
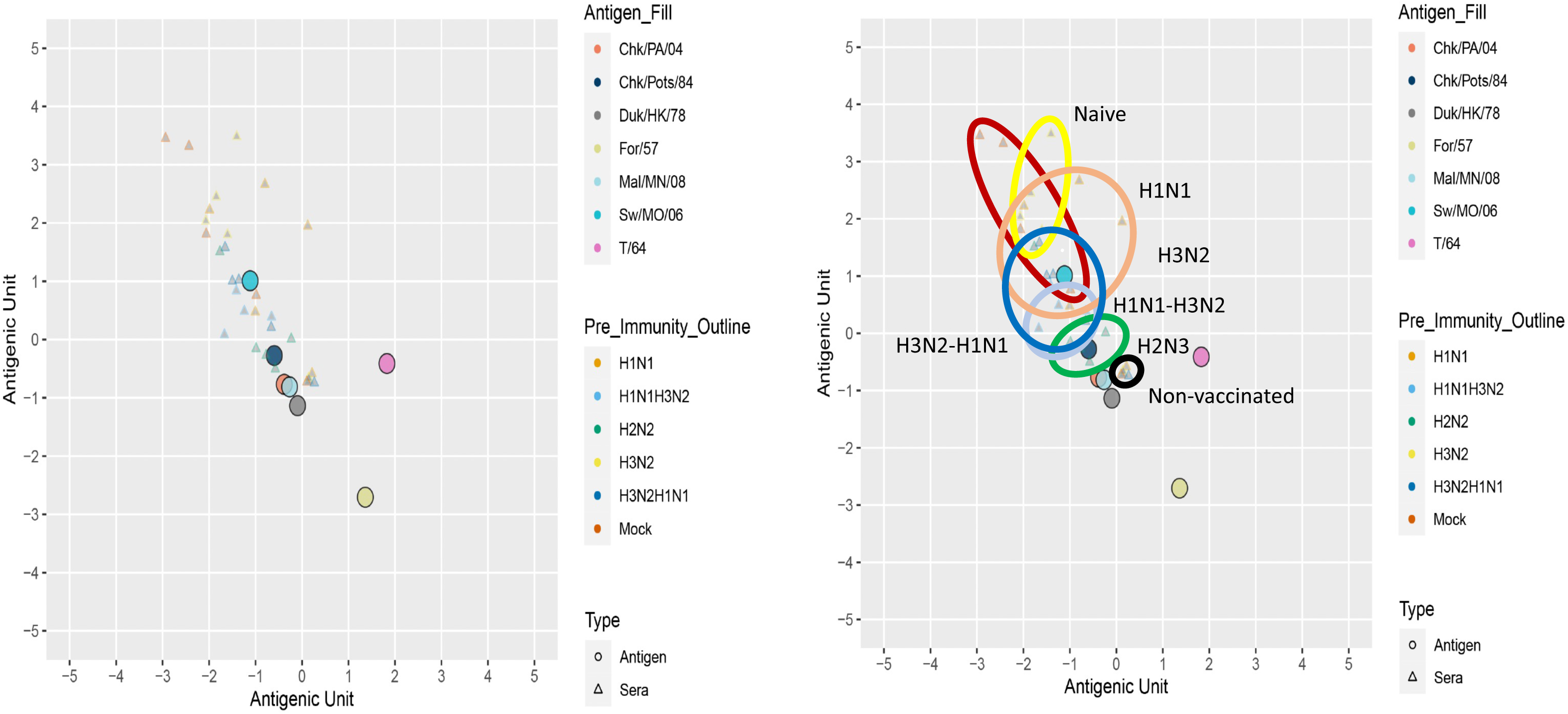
Ferret Neutralization Antigenic Cartography Maps After Boost Vaccination. (A) Antigenic maps were generated for each ferret preimmune group in HAI assays. Antigens are represented as circles and sera from the ferrets are represented by triangles. (B) The sera from each influenza preimmune groups are grouped within each colored circle for clarity.

### Antigenic variations in Human Subjects

Sera from human subjects was evaluated for responses to H2Nx influenza viruses. Serum samples were obtained with informed, written consent in 2016 from participants living in the Athens, GA, USA area. Detailed information about the human study has been previously published and the IRB information is below [9]. Serum was obtained from each participant prior to receiving the seasonal influenza vaccination (split inactivated, tetravalent vaccine, Sanofi-Pasteur, Swiftwater, PA, USA). We used this pre-vaccination serum to evaluate H2 influenza virus immunity. Study participants were divided into four age groups with birth years as follows: 1934-1951, 1952-1966, 1967-1981 and 1982-1996, noting that those born between 1934 and 1966 were likely exposed to H2 strains, while those born after 1967 were likely not exposed. The oldest two age groups (1934-1951 and 1952-1966) had significantly higher HAI titers against thirteen of the sixteen H2 VLPs than serum from participants in the youngest two age groups (1967-1981 and 1982-1996) (Table 4). The oldest two age groups (1934-1951 and 1952-1966) had significantly higher HAI titers than the 1967-1981 age group to the Duk/HK/78 virus, but there was no significant difference in HAI titers against the 1982-1996 age group (Table 4). There were no significant differences in HAI titers between participants in any of the age groups against either the Musk/Rus/14 or the Chk/Pots/84 VLPs (Table 4).

**Table 4:**
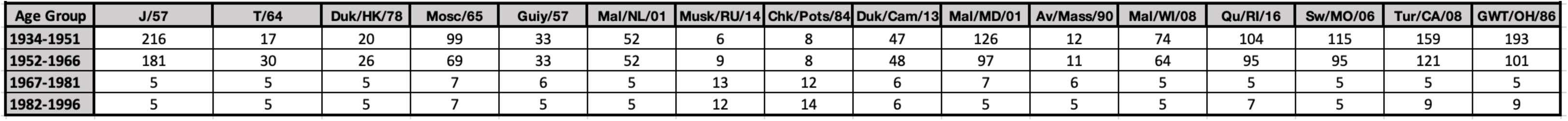
Average Human HAI Titer.

The antigenic cartography for the HAI titers indicated that most of the antigens clustered together near the center of the graph with the Musk/Rus/14 and Chk/Pots/84 antigens being slightly separated from the other antigens (Fig 8). The sera of the 1934– 1951 participants clustered relatively close to each other and several antigenic units away from the 1967-1981 and 1982–1996 birth years age groups (Fig 8). The sera from the 1967-1981 and 1982–1966 groups overlapped but had high variability between participants (Fig 8). The serum collected from participants in the 1952–1966 group had high variation and were several antigenic units apart (Fig 8). This likely reflects that these participants were born both before and after H2N2 entered the human population (Fig 8). The participants born in the mid-1960s also may not have been exposed to H2N2 influenza viruses before these viruses stopped circulating in the human population.

**Figure 8.**
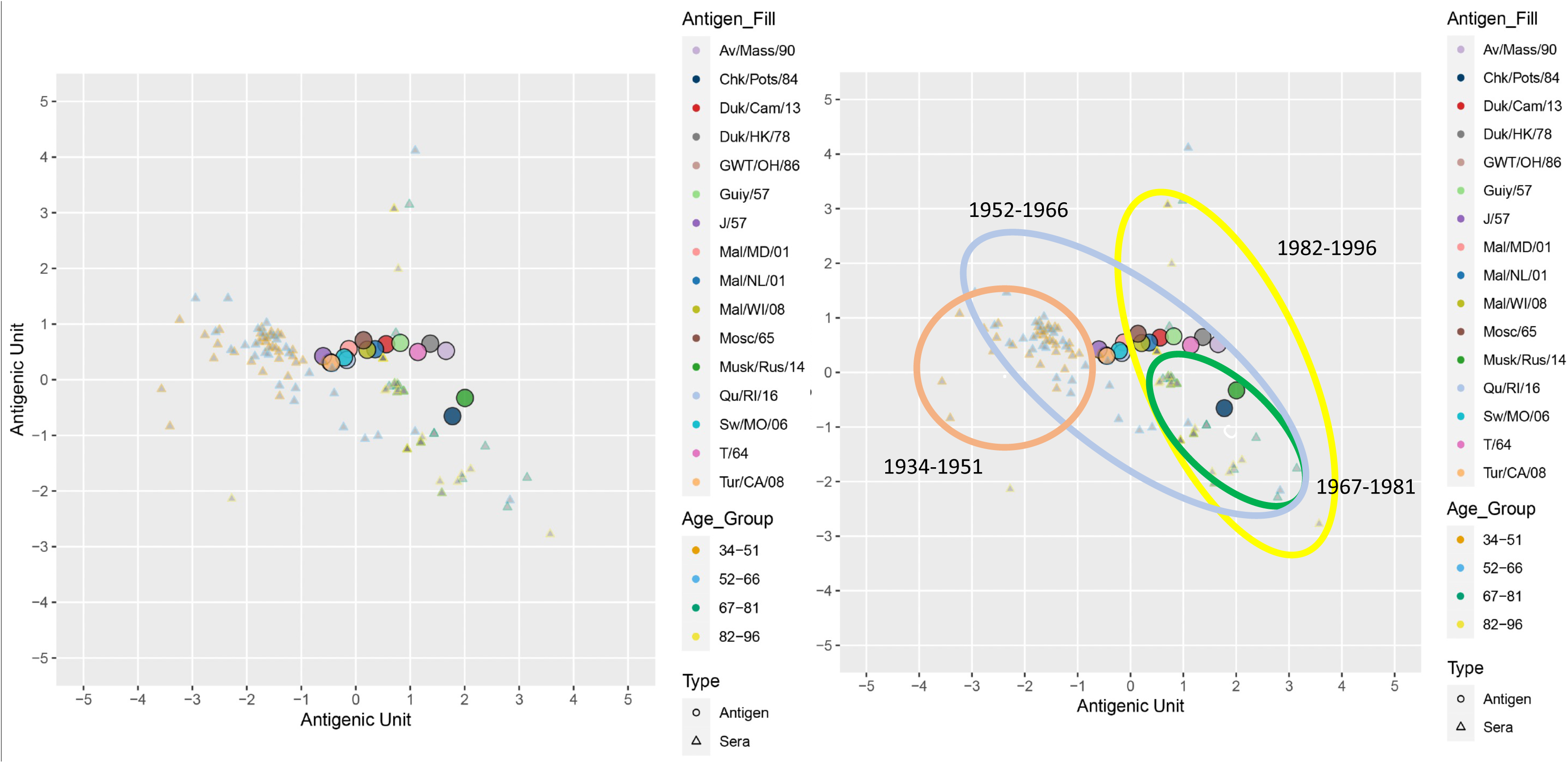
Human HAI Antigenic Cartography Maps. (A) Antigenic maps were generated using a cohort of sera from human subjects in HAI assays. Antigens are represented as circles and sera from the humans are represented by triangles and divided by birth year ranges. (B) The sera from human subjects is divided into groups by years of birth and are grouped within each colored circle for clarity.

To differentiate the sera from individuals born between 1952-1966, we generated antigenic maps of just these individuals (Fig S1). Most of the HA antigens clustered at the center of the map similarly to the map of the entire population (Fig S1). The Chk/Pots/84 and Musk/Rus/14 are divergent from the other antigens that have similarly to the map of the entire population. The T/64, Av/Mass/90 and Duk/HK/78 HA antigens also separated from the other HA antigens (Fig S1). Most of the sera clustered near the center of the map with a few serum samples being highly divergent (>2 antigenic units) (Fig S1).

For neutralization titers, participants in the two oldest age groups had high neutralization titers (>1:400) to Sw/MO/06 and intermediate responses against the other H2 strains, potentially reflecting H1 immune breadth against H2 strains and the circulation of H2N2 strains after 1956 (Table 5). The two oldest age groups had, on average, the highest neut titers (>1:100) against the panel of viruses (Table 5). The participants in the 1952-1966 were the only group to have an average neut titer of ≥1:100 for more than one virus (Table 5). On average, participants in the 1967-1981 age group had reduced/low level H2 titers across the panel. Sw/MO/06 was also the only virus that the 1967-1981 age group had which demonstrated neutralization titers >1:25. (Table 5). As expected, participants in the youngest age group had low neutralization titers (<1:10) against all viruses in the panel (Table 5).

**Table 5:** Average of Human Neutralization Titers.

| Table 5: Average of Human Neutralization Titers |  |  |  |  |  |  |  |  |
| --- | --- | --- | --- | --- | --- | --- | --- | --- |
| Age Groups | Chk/Pots/84 | Chk/PA/04 | Form/57 | T/64 | Duk/HK/78 | Sw/MO/06 | Mal/MN/08 | Average |
| 1934-1951 | 36 | 15 | 73 | 36 | 41 | 444 | 72 | 103 |
| 1952-1966 | 68 | 33 | 36 | 73 | 84 | 424 | 106 | 118 |
| 1967-1981 | 8 | 20 | 19 | 24 | 5 | 83 | 11 | 24 |
| 1982-1996 | 5 | 7 | 5 | 8 | 7 | 6 | 5 | 6 |
| Average | 29 | 19 | 33 | 35 | 34 | 239 | 48 | 63 |

The antigenic cartography for the neutralization assays reveals that all of the antigens except Sw/MO/06 and For/57 cluster together (Fig 9). Sw/MO/06 is at the center of the map, while For/57 is several antigenic distance units from all of the other HA antigens as well as all of the sera (Fig 9). Serum samples from individuals born between

**Figure 9.**
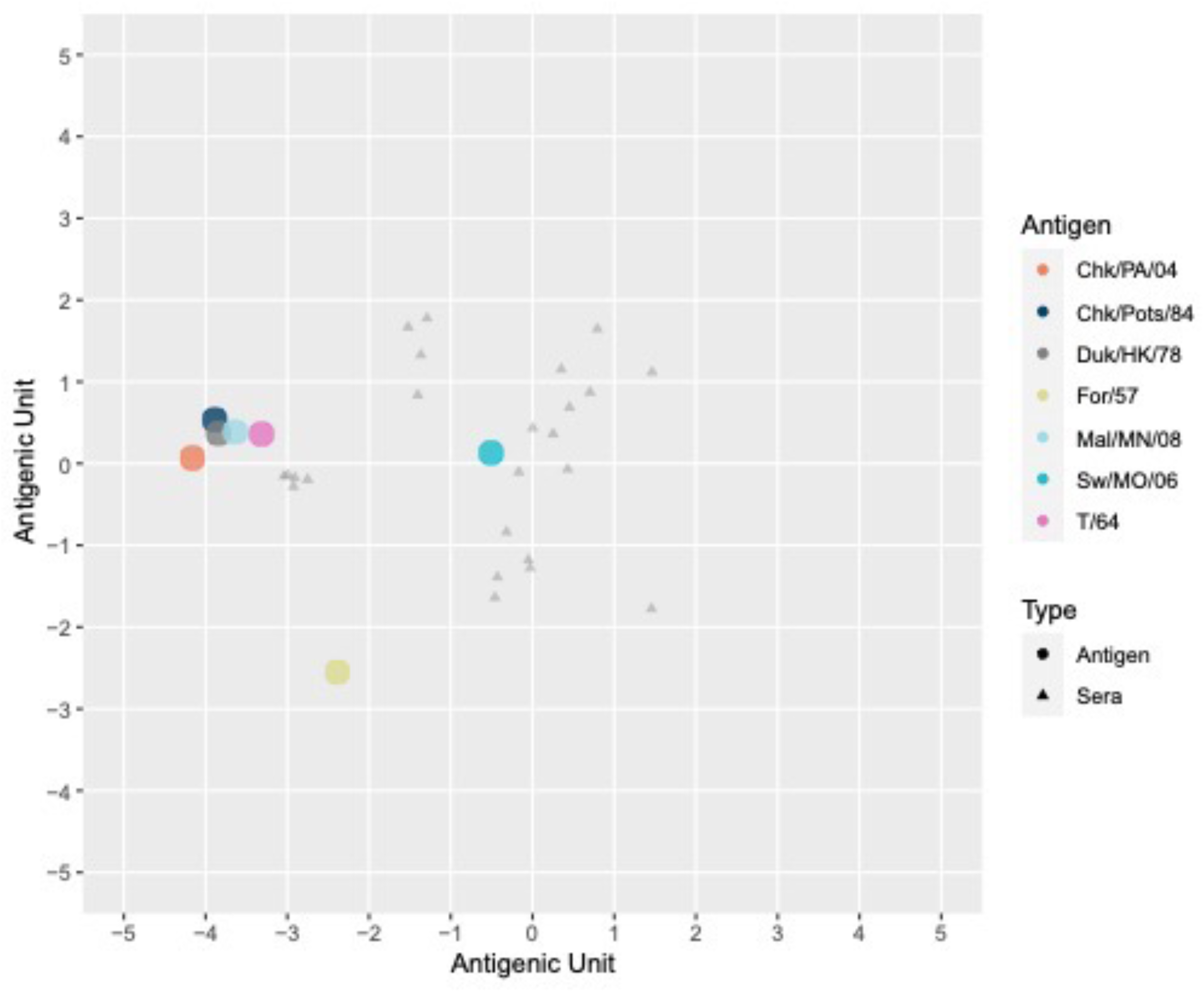
Human Neutralization Antigenic Cartography Maps. Antigenic maps were generated using a cohort of sera from human subjects in neutralization assays. Antigens are represented as circles and sera from the humans are represented by triangles and divided by birth year ranges. The Sw/MO/06 antigen is nearest to the sera with For/57 being divergent from the other antigens and the remaining five antigens grouping together apart from the majority of the sera.

1937 and 1958 cluster together near the center-right of the map [(-0.5,0) and (2,2)] (Fig 9). Serum from participants born between 1942 and 1966 cluster together in the bottom center portion of the map [(-0.5,-1) and (2,2)] (Fig 9). Participants in both of these groups likely contain individuals infected one or more times with H2N2 influenza viruses. Serum samples from participants born between 1947 and 1954 cluster together in the upper left portion of the map [(-1,0) and (-2,2)] (Fig 9). These are likely participants who were initially imprinted with H1N1 and then coinfected with an H2 influenza virus strain. Finally, serum samples collected from participants born between 1966 and 1996 cluster together on the left side of the map [(-2,-1) and (-3.5,-1)] (Fig 9). These are likely participants who were never infected with H2N2 influenza viruses.

## Discussion

Using antigenic cartography to map the antigenic distances between H2 influenza virus strains, we re-analyzed both HAI and neut data in animals and in humans to evaluate the impact of variable single and multiple H1 and H3 infection preexposure histories on H2 vaccine performance. Under controlled experimental conditions, these studies collectively reveal the stark differences in H2N2 antigenic recognition between different animal species and humans, depending upon exposure history. In particular, a primary H1N1 infection or H1N1 + H3N2 infection elicits imprinted responses the effectively promote robust immune titers following H2 vaccination, while H3N2 primary infections (regardless of H1N1 secondary infection) attenuate H2 vaccine responses. Obviously, these conclusions are based on single exposures of these strains, and may underrepresent the impact of multiple, sequential infections on H2 vaccine performance.

Ferrets, pre-exposed to either H1N1 or H3N2 alone or after sequential infection with both viruses elicited little if any cross-neutralization titers directed against H2N2 influenza viruses isolated over a 50-year period. After rH2 vaccination, H2N3 preimmune ferrets had neutralization titers that were significantly increased following vaccination with homologous H2 HA antigens against all H2 subvariants in the panel, although some strains were significantly more resistant to neutralization than others. In contrast, ferrets pre-exposed to H3N2 influenza viruses either alone or prior to H1N1 influenza virus infection had significantly reduced the titers of all H2 vaccines. Importantly, these attenuated responses can be modulated in part, using the Z1 COBRA immunogen which elicited superior H2 responses as compared with the other H2 HA antigens tested.

One likely interpretation of these data is that H2 HA vaccines may be less effective in preimmune, adult populations, depending on exposure history and the sequence of H1N1 and H3N2 infection/vaccine history as well as host genetic variation. The data also suggests that naïve infants will respond differently than adult populations after prime vaccination or infection due to the absence of memory B cells. Additional studies are needed to evaluate the impact that vaccination following initial influenza exposure has on future influenza vaccinations, as opposed to infections. Moreover, these studies reveal that factors such as preimmune status, strains used in viral panel, and host genetic background can all obfuscate the results of vaccine studies and should be carefully considered in future clinical studies.

H2 HA vaccine boosters clearly enhanced global H2 neutralization responses across the panel, regardless of vaccine group, with interesting caveats. Overall, the Z1 COBRA HA vaccine was clearly superior to all other formulations as measured by HAI and neut titers. Still, one H2 HA (Musk/RUS/14) was clearly resistant to neutralization suggesting an important role for strain specific variation in altering neutralization performance and potentially leading to breakthrough infections. After the second vaccination, both the H3N2 and H3N2-H1N1 preimmune ferrets had higher H2 HAI titers across the panel, although overall, were slightly lower that the titers in H2 preimmune-H2 vaccinated ferrets. The stark differences between the first and second vaccinations in the HAI assay suggest the need for multiple vaccinations in the event of future H2Nx influenza pandemic. This would be similar to the multiple vaccinations required to generate robust antibodies against the SARS-CoV-2 virus, since people had no pre-existing immunity to this virus that cause COVID-19. Most people under the age of 60 do not have pre-existing neutralizing antibodies against H2Nx influenza viruses [9].

The strains Musk/Rus/14, Duk/HK/78 and Duk/Cam/13 were the most divergent across the various ferret preimmune groups. When comparing the sequences of these three HA proteins, there is no single mutation shared between the three HA antigens that is different from the other HA proteins in the panel. This suggests that multiple mutations, working alone or in concert, have the potential to alter the ability of the HA vaccine to elicit broad neutralization activity. Indeed, multiple amino acids substitutions are responsible for antigenic changes in human H2N2 influenza virus diversity [2, 14]. The ferret neut data from each group highly overlapped mainly due to the high variation in the mock vaccinated ferrets. The T/64 and For/57 HA antigens are the most divergent from the other HA antigens, reflecting the low neutralizing titers across all ferret groups.

The antigenic cartography of H2 influenza HA in humans also identified patterns that can be further analyzed for a better understanding of viral variants and their antigenic differences over time. Most humans are infected with and likely imprint on the first infecting influenza virus strain in four years of birth [6, 7]. These responses will likely vary for individuals across different age groups as drift alters the prevalent strains (or vaccines) over time. Such trends in the antigenic cartography maps were highlighted in four different age groups: 1934-1951, 1952-1966, 1967-1981, 1982-1996. The oldest demographic and the second oldest demographics had nearly the same antigen recognition profile that was focused mainly on H2N2 Sw/MO/06 and low neutralizing titers (<1:80) to the majority of the other H2Nx influenza viruses in the panel. A finding that likely points to a previous exposure to H2N2 influenza viruses. However, the overall lack of broadly neutralizing H2 influenza immunity suggests that H2-specific immunity in older individuals has waned considerably over the next 50+ years. In contrast, the younger cohorts only neutralized three H2 HA antigens due to one participant sera that recognized J/57, GWT/OH/86, and Tur/CA/08. However, sera from more participants within this age range are needed to validate these outcomes. This large separation in our HAI human map likely stems from the difference in immunological imprinting that occurred during different periods of history and suggests head specific antibody recognition correlated to immunity. Participant born between 1952-1966 had the broadest range of antigen recognition, including all the H2 HA antigens with a trend that included Mal/WI/08, Mosc/65, Mal/MD/01, Qu/RI/16, and Sw/MO/06.

This study also utilized data generated by Linster *et al.* [2], who identified significant antigenic sites in HA by rescuing human derived H2N2 influenza viruses with single amino acid mutations. These mutant HA sequences were compared on the antigenic maps generated using HAI and neutralization titers. Some mutations found by Linster *et al.* likely explain the divergence of antigens, such as Musk/Rus/14 and T/64 [2]. While other divergent HA antigens, such as Duk/HK/78 did not have significant amino acid changes in the sites identified by Linster *et al.* The data from this study suggests that additional significant antigenic sites at positions 140 and 151 may contribute to the divergence of Musk/Rus/14, T/64 and Duk/HK/78 either singly or in combination, although, there are numerous sites where Duk/HK/78 encodes unique amino acids. The high number of unique mutations in Duk/HK/78 would require extensive additional studies to ascertain which sites are causing antigenic divergence. These sites may not have been identified previously because the human derived H2N2 viruses did not contain mutations at either of these amino acid locations. We attempted to elevate the significance of both antigenic sites, but we were unable to show statistically significant HAI changes for either. Additional studies are needed to investigate the antigenic significance of each mutation in Musk/Rus/14 HA, both individually or in combination.

The antigenic maps compared data across species and evaluated antigenic sites. The extensive data sets generated in this study will provide valuable information for future H2 influenza virus immunity and the generation of H2 influenza vaccines. Additionally, using antigenic cartography to identify divergent strains for further evaluation can be applied to other pathogens.

## Materials and Methods

### COBRA HA antigen design

The design of each COBRA vaccine has been previously published in detail [13]. Briefly, amino acid sequences from wild-type (WT) H2Nx strains were downloaded from the Global Initiative on Sharing All Influenza Data (GISAID). The original amino acid sequences were grouped together to create primary consensus sequences. This layering consensus method was continued until final consensus sequences were obtained. These COBRA amino acid sequences were then reverse translated into nucleotide sequences and codon optimized for mammalian expression to create VLPs.

In response to the intensive task of applying the Computationally Optimized Broadly Reactive Antigen (COBRA) method within Geneious Prime, a program was designed leveraging the Geneious Prime Application Programming Interface and Java to streamline the process by automating several key steps. First, the program automates the alignment of amino acid sequences, ensuring that despite sequence diversity, all amino acids at corresponding positions are oriented. Next, specific regions of proteins, typically variable ones, are automatically selected, truncated, and followed by a second round of alignment. Subsequently, the program groups the realigned sequences based on a predetermined similarity threshold, effectively clustering sequences that share resemblance to a reference sequence. Within each group, it generates a consensus sequence representing the most common amino acid at each position across the sequences. The various groupings of consensus sequences establish a primary layer, which serves as the foundation for further refinement.

Following the creation of the primary layer, the program iteratively groups consensus sequences from the previous layer into a new layer, continuing this process until convergence to a single sequence suitable for vaccine development is obtained. Through automation, the program has significantly expedited the implementation of the COBRA method within Geneious Prime, streamlining the process of identifying potential vaccine candidates from diverse amino acid sequences while minimizing human error.

### Ferret vaccinations

Fitch ferrets (Mustela putorius furo, spayed, female, 6 to 12 months of age) were purchased certified influenza-free and de-scented from Triple F Farms (Sayre, PA, USA). Ferrets were pair housed in stainless steel cages (Shor-Line, Kansas City, KS) containing Sani-Chips laboratory animal bedding (P. J. Murphy Forest Products, Montville, NJ). Ferrets were provided with Teklad Global Ferret Diet (Harlan Teklad, Madison, WI) and freshwater ad libitum. The University of Georgia Institutional Animal Care and Use Committee approved all experiments, which were conducted in accordance with the National Research Council’s Guide for the Care and Use of Laboratory Animals, the Animal Welfare Act, and the CDC/NIH’s Biosafety in Microbiological and Biomedical Laboratories guide. Ferrets (n=20) were infected with H1N1, or H3N2 seasonal influenza viruses or H2N2 avian influenza viruses in different orders before vaccination. These influenza viruses included the H1N1 influenza viruses Singapore/6/1986 (Sing/86) and California/07/2009 (CA/09), the H3N2 influenza viruses Sichuan/2/1987 (Sich/87) or Panama/2007/1999 (Pan/99), and the H2N2 avian influenza viruses Chk/PA/04 or Qu/RI/16, all at an infectious dose of 1e+6 PFU in 1mL intranasally. For the ferrets with multiple preimmune infections, ferrets were left for 60 days between each infection and before the first vaccination.

After the establishment of preimmunity by viral infection, 60 days elapsed before ferrets were vaccinated with recombinant hemagglutinin (rHA) twice with 4 weeks between vaccinations. The ferrets were vaccinated with a 1:1 ratio (500ml total volume) of rHA diluted with phosphate-buffered saline (PBS) (15.0ug rHA/ferret) and the emulsified oil-water adjuvant Addavax (InvivoGen, San Diego, CA, USA). The mock-vaccinated groups received only PBS and Addavax adjuvant at a 1:1 ratio (500ml total volume) with no rHA. Each vaccination was given intramuscularly. Before vaccinations and 2 weeks after each of the vaccinations, ferrets were bled and serum was isolated from each of the samples. The blood was harvested from all anesthetized ferrets via the anterior vena cava at days 0, 14, and 42. Blood samples were incubated at room temperature for 1 hr prior to centrifugation at 6,000 rpm for 10 min. The separated serum was removed and frozen at -20°C.

### Mouse vaccinations

BALB/c mice (females, 6 to 8 weeks old) were purchased from Jackson Laboratory (Bar Harbor, ME, USA) (Stock #000671). The mice were housed in microisolator units and were given both water and food ad libitum. The mice were vaccinated with a 1:1 ratio (50μL total volume) of rHA diluted with phosphate-buffered saline (PBS) (3.0μg rHA/mouse) and the emulsified oil-water adjuvant, Addavax (InvivoGen, San Diego, CA, USA). The mice were boosted with the same vaccine formulation with the same dosage at 4 weeks post-initial vaccination. Blood samples were obtained from the mice via cheek bleeds fourteen to eighteen days following the second vaccination. Blood samples were collected in 1.5 mL microcentrifuge tubes. The blood samples were incubated at room temperature for one hour and then centrifuged at 6000 rpm for ten minutes. Serum samples were transferred to new 1.5 mL microcentrifuge tubes and stored at -20°C.

### Animal Ethics

Animal were cared for under the University of Georgia (UGA) Research Animal Resources guidelines for laboratory animals. All procedures were reviewed and approved by the Institutional Animal Care and Use Committee (IACUC).

### Human Participants and Vaccinations

Participants ranging between the ages of 19-82 years old consented to the study and were enrolled in Athens, GA, USA. Various factors were utilized to determine the participants’ eligibility. Those who had not yet received the seasonal influenza virus vaccine at the time of enrollment, the beginning of September 2016, were still included in the study (University of Georgia, IRB# STUDY00003773). Influenza strains included in the vaccine were based upon the WHO recommendations for the Northern hemisphere: (A/California/7/2009-H1N1), (A/Hong Kong/4801/2014H3N2), (B/Phuket/3073/2013-Yamagata-lineage), (B/Brisbane/60/2008-Victoria-lineage). Participants received vaccinations from September 2016 to December 2016. Participants were vaccinated with the standard dose (15 μg/antigen) split-virion (IIV) version of licensed Fluzone (Sanofi Pasteur) influenza virus vaccine. 148 participants were enrolled in the study. Approximately 80 mL of blood was collected from each participant before vaccination (D0), 7 days post infection (D7), and 21 days post-infection (D21). Sera and peripheral blood mononuclear cells (PBMCs) were isolated from the blood samples. Sera was collected in Vacutainer serum separation tubes (SST) tubes (BD Biosciences) and processed within 48 hours, aliquoted and stored at -20 °C. PBMCs were collected in Vacutainer cell preparation tubes (CPT) tubes (BD Biosciences) at D0, D7 and D21. PBMCs samples suspended in DMSO and FBS and were stored in liquid nitrogen.

### Viruses, rHA antigens and VLPs

The following were obtained from either the United States Department of Agriculture’s (USDA) Diagnostic Virology Laboratory (DVL) in Ames, Iowa; BEI resources; or provided by the laboratory of Dr. S. Mark Tompkins in Athens, GA: A/Chicken/Potsdam/4705/1984 (Chk/Pots/84), A/Chicken/PA/298101-4/2004 (Chk/PA/04), A/Duck/Hong Kong/273/1978 (Duk/HK/78), A/Mallard/Minnesota/AI08-3437/2008 (Mal/MN/08), A/Swine/Missouri/4296424/2006 (Sw/MO/06), A/Formosa/313/1957 (For/57), A/Japan/305/1957 (J/57), and A/Taiwan/1/1964 (Tw/64). The following H1N1 influenza viruses used in the study were provided by either the Centers for Disease Control and Prevention (CDC) or Virapur LLC: A/Singapore/6/1986 (Sing/86), A/California/07/2009 (CA/09; pandemic). Passaged using embryonated chicken eggs, each virus was harvested from the eggs and aliquoted into tubes which were stored at -80 °C. Each virus was tittered using a standard influenza plaque assay.

Recombinant HA (rHA) proteins were produced using the pcDNA 3.1+ plasmid. Each HA gene was truncated by removing the transmembrane (TM) domain and the cytoplasmic tail at the 3’ end of the gene. The TM domain was determined using the TMHMM Server v. 2.0 website: http://www.cbs.dtu.dk/services/TMHMM/. The HA gene was truncated at the first amino acid prior to the TM domain. A fold-on domain from T4 bacteriophage, an Avitag and a 6X histidine tag totaling 477 nucleotides were added to the 3’ end of the HA gene. The pcDNA 3.1+ vectors were then transfected individually into HEK293T suspension cells using ExpiFectamine 293 transfection reagent following manufacturer’s specifications (ThermoFisher Scientific). The supernatants were then harvested from the transfected HEK293T cells. rHAs were purified from the supernatant using a nickel-agarose column. The rHAs were then eluted from the column using imidazole. After elution, proteins were quantified using bicinchoninic assay (BCA) and stored at -80 °C.

For the virus-like particle (VLP) production, human endothelial kidney 293T (HEK-293T) cells (1 x 10^6^) were transiently transfected for the creation of mammalian virus-like particles (VLPs). DNA of each of the three pcDNA 3.1+ mammalian expression vectors expressing the influenza neuraminidase (A/South Carolina/1/1918; H1N1), the HIV p55 Gag sequence, and one of the various H2 wild-type HA proteins was added in a 1:2:1 ratio with a final DNA concentration of 1μg. Following 72 h of incubation at 37°C, supernatants from transiently transfected cells were collected, centrifuged to remove cellular debris, and filtered through a 0.22 μm pore membrane. VLPs were purified and sedimented by ultracentrifugation on a 20% glycerol cushion at 23,500g for 4h at 4°C. VLPs were resuspended in phosphate buffered saline (PBS), and total protein concentration was determined with the Micro BCA Protein Assay Reagent kit (Pierce Biotechnology, Rockford, IL, USA). Hemagglutination activity of each preparation of VLP was determined by serially diluting volumes of VLPs and adding equal volume of 0.8% turkey red blood cells (RBCs) (Lampire Biologicals, Pipersville, PA, USA) suspended in PBS to a V-bottom 96-well plate with a 30 min incubation at room temperature (RT). Prepared RBCs were stored at 4°C and used within 72 h. The highest dilution of VLP with full agglutination of RBCs was considered the endpoint HA titer. The HA sequences used for VLPs were Mal/NL/01, Chk/Pots/84, Muskrat/Russia/63/2014 (Musk/Rus/14), Duck/Cambodia/419W12M3/2013 (Duk/Cam/13), J/57, Moscow/1019/1965 (Mosc/65), T/64, Duk/HK/78, Mal/WI/08, Sw/MO/06, Quail/Rhode Island/16–018622-1/2016 (Qu/RI/16), Turkey/California/1797/2008 (Tur/CA/08).

### Hemagglutination-inhibition assay

Hemagglutination inhibition (HAI) assays were used to quantify receptor-binding HA-specific antibodies through measuring the inhibition in the agglutination of turkey RBCs. Prior to being tested, the sera were treated with receptor-destroying enzyme (RDE) (Denka Seiken, Co., Japan) to inactivate nonspecific inhibitors. Three parts RDE was added to one-part sera and incubated overnight at 37 °C. RDE was subsequently inactivated by incubating the serum-RDE mixture at 56 °C for approximately 45 min. After the incubation period, six parts PBS was added to the RDE-treated sera. The RDE-treated sera were two-fold serially diluted in V-bottom microtiter plates. An equal volume of each VLP was adjusted to approximately 8 hemagglutination units (HAU)/25 μL and added to each well of the V-bottom microtiter plates. The plates were covered and incubated at RT for 20 min before adding 50 μL of RBCs which were allowed to settle for 30 min at RT. The HAI titer was determined by the reciprocal dilution of the last well that contained non-agglutinated RBCs.

### Neutralization assays

Neutralization assays were used to identify the presence of virus-specific neutralizing antibodies. Antibodies were diluted in ½-log increments with serum-free media and incubated with 100 times TCID50 for 1 h. The antibody-virus mixture was then added to the incomplete (FBS-free) DMEM washed MDCK cells in the 96-well plate. After 2 h, the MDCK cells were washed with incomplete DMEM. Approximately 200 μL of DMEM with P/S and TPCK was added to each of the 96 wells. The cell monolayers were checked daily for cytopathic effect (CPE). CPE was defined as > 10% CPE of cells per well. After 3-4 days, media in each well was removed and the MDCK cells were fixed with 10% buffered formalin. MDCK cells were stained using 1% crystal violet (Sigma).

### Study Approval

Study procedures, informed consent, and data collection documents were reviewed and approved by the IRB of the University of Georgia. Subjects were recruited at a medical facility in Athens, GA, USA and enrolled with written, informed consent. Exclusion criteria included documented contraindications to Guillain-Barré syndrome, dementia or Alzheimer’s disease, allergies to eggs or egg products, estimated life expectancy <2 years, medical treatment causing or diagnosis of an immunocompromising condition, or concurrent participation in another influenza vaccine research study. Participants were not monitored for influenza virus infection during the time of their participation. Although influenza virus did not circulate widely in the community during this time, participants were asked during each visit if they had experienced flu-like symptoms, and those who did were excluded from the study.

### Antigenic Cartography

The antigenic cartography analyses were carried out in R version 4.1.3 (https://www.R-project.org/) with the antigenic and sera coordinates calculated through the racmacs package (https://github.com/acorg/Racmacs) and the graphic displays done with ggplot2.<u>^94^</u> The samples were grouped by either vaccine, preimmunity, or age. The coordinates of the sera and antigens were calculated with a run of 1000 optimizations to minimize the difference between the n-dimensional Euclidean distances between points, and the two-dimensional distances on the final map.

### Statistics

Statistical significance for assays was defined as a p-value of less than 0.05. Limit of detection for HAIs is 1:10. 1:5 was used for statistical analysis. The HAI titers were transformed by log2 for analysis and graphing for both paired *t*-test and one-way ANOVA analysis. Geometric mean titers were calculated for neutralization assays, but the Log 10 titers were used for both paired *t*-test and ANOVA analysis. All error bars on the graphs represent standard mean error.

## Author Contributions

Z.B.R., conceptualization, formal analysis, methodology, writing; C.N., figure generation; M.R.C., formal analysis, writing; J.L.P., writing; O.C.R., formal analysis, writing; E.O., writing; T.M.R., conceptualization, formal analysis, funding acquisition, methodology, writing. R.S.B., conceptualization, formal analysis, methodology, writing.

## Institutional Review Board Statement

The study was conducted in accordance with the Declaration of Helsinki and approved by the Ethics Committee of WIRB and University of Georgia IRB (Study #00003773). The studies were conducted in accordance with the local legislation and institutional requirements. The participants provided their written informed consent to participate in this study.

## Informed Consent Statement

Informed consent was obtained from all subjects involved in the study.

## Data Availability Statement

All of the raw data used for this study has been previously published [8, 9, 13]. Data from these studies can also be provided upon request.

## Acknowledgments

The authors acknowledge the UGA CIVIC Flu Vaccine teams for sample collections blood and saliva processing and technical assistance. We would also like to thank all participants enrolled in the study, as well as Dr. Brad Phillips, Kimberly Schmitz, and the entire staff at the University of Georgia Clinical and Translational Research Unit (CTRU) for assistance in collecting samples in the influenza vaccine program. The CTRU was supported by the National Center for Advancing Translational Sciences of the National Institutes of Health under Award Number UL1TR002378.

---

**Figure S1.**
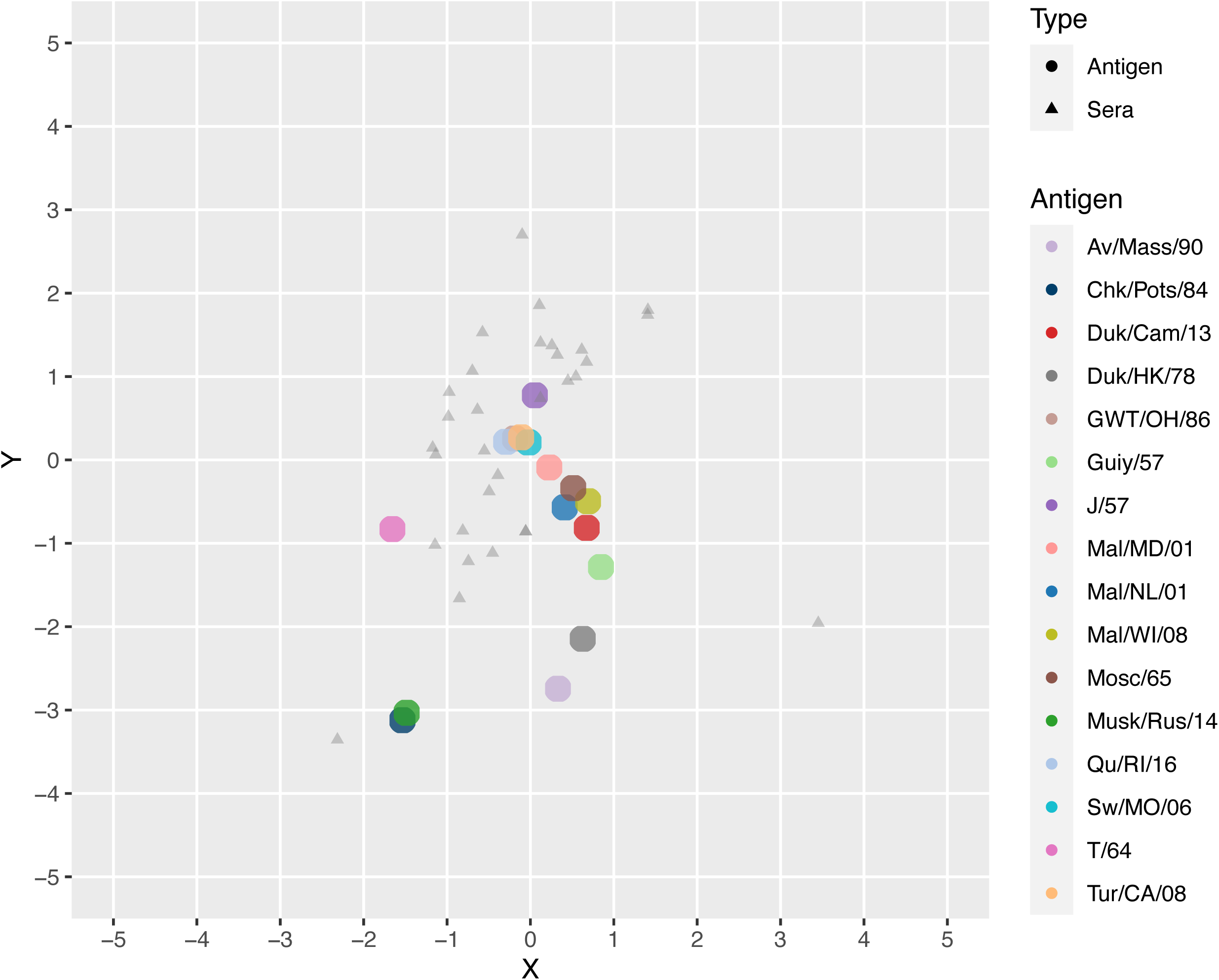
Human HAI Antigenic Cartography Maps for 1952-1966 Age Group. Antigenic maps were generated using a cohort of sera from human subjects born between 1952 and 1966 in HAI assay. These individuals were born either just before or during H2N2 circulation in humans. Antigens are represented as circles and sera from the humans are represented by triangles. The sera largely groups together around the majority of the antigens. The Chk/Pots/84 and Musk/Rus/14 antigens are the most divergent from the other antigens.

**Table S1:** Sites from Linster et al.

| Table S1: Sites from Linster et al. |  |  |  |  |  |  |  |
| --- | --- | --- | --- | --- | --- | --- | --- |
| Strain | 126 | 128 | 132 | 139 | 154 | 184 | 188 |
| Chicken/Potsdam | T | T | Q | N | S | T | T |
| Duck/Cambodia | T | T | R | N | S | A | A |
| Duck/Hong Kong | T | T | R | N | S | T | T |
| Japan/57 | T | T | R | N | S | T | T |
| Mallard/netherlands | T | T | R | N | S | A | A |
| Mallard/Wisconsin | T | T | R | N | S | A | T |
| Moscow/1965 | T | T | R | N | S | T | T |
| Muskrat/Russia/14 | T | T | Q | K | S | A | A |
| Quail/RI/16 | T | T | R | N | S | A | A |
| Swine/MO/06 | T | T | R | N | S | A | T |
| T/64 | T | T | M | K | T | A | A |
| Turkey/CA/08 | T | T | R | N | S | A | T |
| Z1 | T | T | R | N | S | A | T |
| Z3 | T | T | R | N | S | A | T |
| Z5 | T | T | R | N | S | T | T |
| Z7 | T | T | R | N | S | A | T |

**Table S2:** Other potential sites.

| Table S2: Other potential sites |  |  |
| --- | --- | --- |
| Strain | 140 | 151 |
| Chicken/Potsdam | P | K |
| Duck/Cambodia | P | K |
| Duck/Hong Kong | P | T |
| Japan/57 | P | K |
| Mallard/netherlands | P | K |
| Mallard/Wisconsin | P | K |
| Moscow/1965 | P | K |
| Muskrat/Russia/14 | S | K |
| Quail/RI/16 | P | K |
| Swine/MO/06 | P | K |
| T/64 | P | E |
| Turkey/CA/08 | P | K |
| Z1 | P | K |
| Z3 | P | K |
| Z5 | P | K |
| Z7 | P | K |

